# Agent-based modeling of cellular dynamics in adoptive cell therapy

**DOI:** 10.1101/2025.02.17.638701

**Authors:** Yujia Wang, Stefano Casarin, May Daher, Vakul Mohanty, Merve Dede, Mayra Shanley, Rafet Başar, Katayoun Rezvani, Ken Chen

## Abstract

Adoptive cell therapies (ACT) leverage tumor-immune interactions to cure cancer. Despite promising phase I/II clinical trials of chimeric-antigen-receptor natural killer (CAR-NK) cell therapies, molecular mechanisms and cellular properties required to achieve clinical benefits in broad cancer spectra remain underexplored. While *in vitro* and *in vivo* experiments are required in this endeavor, they are typically expensive, laborious, and limited to targeted investigations. Here, we present ABMACT (Agent-Based Model for Adoptive Cell Therapy), an *in silico* approach employing agent-based models (ABM) to simulate the continuous course and dynamics of an evolving tumor-immune ecosystem, consisting of heterogeneous “virtual cells” created based on knowledge and omics data observed in experiments and patients. Applying ABMACT in multiple therapeutic context indicates that to achieve optimal ACT efficacy, it is key to enhance immune cellular proliferation, cytotoxicity, and serial killing capacity. With ABMACT, *in silico* trials can be performed systematically to inform ACT product development and predict optimal treatment strategies.

## 1. Main

Adoptive cell therapies (ACT) have shown substantial progress in combating cancer and other diseases^1–9^. By administrating lymphocytes with intrinsic or engineered antitumoral capabilities, ACTs harness the power of the immune system to eliminate tumors^9^. Recent Chimeric-antigen-receptor natural killer (CAR-NK) cell therapies have been identified as a promising alternative to CAR-T cell therapies, given their ability to address multiple limitations of CAR-T therapies^2,10–15^. As a low-cost, low-toxicity, off-the-shelf solution, various CAR-NK cell therapy products have burgeoned to deliver benefits in a broad spectrum of clinical applications^12,13^. CAR extracellular domain engineering provides specificity for anti-tumor effects, such as anti-CD123 targeting acute myeloid leukemia cells^16^, anti-CD19 targeting B cell lymphoma^17^, and anti-CD70 for hematological malignancies and solid tumors^18^. Various genetic engineering approaches can augment CAR-NK cell functions: armoring NK cells with co-stimulatory cytokine vectors such as IL-15^19^, IL-21^20^, and STING agonist^21^ can boost NK cell proliferation and killing capability, while CRISPR editing such as *CISH* deletion^22^ increases NK cell metabolic fitness. In the case of cord-blood derived CAR-NK cells, donor characteristics can dictate CAR-NK function and clinical efficacy. Increased inflammation, hypoxia, and cellular stress have been associated with suboptimal cord blood preservation, and CAR-NK cells engineered from these cords have reduced anti-tumor efficacy^5,23^. Despite significant advances, applications of CAR-NK ACT are predominantly in preclinical and phase-I/II trial stages^1,14^, highlighting both the potential and need for further investigation.

As “living drugs”, ACTs present distinct opportunities and challenges in clinical development^24^. ACT achieves treatment responses through interactions between the product and the target cells. The cell products are often genetically modified to boost therapeutic potential. However, both aspects are difficult to assess. While *in vitro* and *in vivo* models have been instrumental in advancing ACT development, they are often costly, labor-intensive, and limited in their ability to replicate the complexity of the interactions between the human immune system and the tumor microenvironment (TME)^25,26^. The TME is characterized by diverse and evolving cell populations, along with a rich and dynamic molecular environment, which cannot be fully recapitulated using traditional cell-line models^26^. Organoids are engineered to replicate the morphology and functions of tissues and organs, yet significant technological challenges remain^27^. As a result, organoid cultures still struggle with repeatability and face challenges in cell maturation and accurately replicating the complexity of native tissue^27^. Although patient-derived xenograft (PDX) animal models aim to reflect TME heterogeneity, they still diverge significantly from human physiology and pharmacology, limiting their translational relevance^25^. Perturbing PDX models is time consuming and expensive, limiting exploration and hypothesis testing. Further, evaluating treatment responses, particularly cellular kinetics and molecular interactions, such as peak concentration and target interaction rates, remains a challenge in these experimental systems due to the difficulties of continuous monitoring^24^. A more comprehensive understanding of multi-scale dynamics at both cellular and sub-cellular level is essential to advance ACT development. This includes capturing the heterogeneous and evolving cell populations, modeling phenotypic and functional states of individual cells, and accounting for their molecular variations. Addressing these challenges requires innovative approaches that combine experimental data with mechanistic and computational modeling, which have the potential to provide deeper insights into complex biological dynamics.

Mathematical and computational models have been utilized to understand complex biological systems, spanning from organisms to molecules^28–33^. Recent advancements in ACTs and the growing availability of data have facilitated their modeling, though further efforts are needed, particularly in emerging areas such as engineered NK cell therapy. Machine learning models such as random forests, support vector machines, and neural networks can learn relationships between entities from data but have limited biological interpretability. This limitation reduces their utility for generating mechanistic insights and translating findings into clinical applications^34^. Sparse data points collected in *in vitro* or *in vivo* experiments also restrict these models’ ability to accurately capture the dynamics of ACT. Mechanistic models such as ordinary differential equations (ODE), partial differential equations (PDE), and stochastic differential equations (SDE) have been developed to describe the kinetics of cell populations and cytokines in the TME^35^. While these approaches offer improved efficiency and mathematical interpretability, they often fall short in capturing molecular and populational heterogeneity, spatial interactions, and stochasticity at the cellular level^36^. Among the vast suite of *in silico* models, Agent-Based Models (ABM)^37,38^, also known as individual-based models, offer the unique advantage of simulating the tumor microenvironment using a bottom-up approach to model individual cell behaviors^28,39^. ABMs represent cells and molecules in the TME as agents and environment attributes, in which the behaviors of each type of agent are coded via simplifications and approximations of biological processes^37,38^. ABM offers several advantages, including temporal prediction capabilities, spatial distribution insights, interpretability, and allow modeling intrinsic stochasticity at multiple scales of the biological systems, overcoming many limitations of other modeling methods like ordinary differential equations and neural networks^28^. Extensive research has demonstrated the potential of using ABM to reproduce the complex interactions in the TME and predict tumor progression or treatment outcomes^36,40–43^. Several studies have attempted to model heterogeneity of cell states by summarizing functional effects of genes^36,39,41,44^. However, these effects are usually binarized as positive and negative regulators of cellular functions without considering their relative importance. Moreover, the application of ABMs to cellular immunotherapies, particularly engineered NK cell therapies, remain unexplored. Several critical questions must be addressed: How can a system of cell agents be designed to sufficiently capture the diversity of cell types and phenotypic transitions in NK-ACT? How can molecular profiles be integrated with cellular functions to model functional heterogeneity? How can knowledge and data be balanced to ensure both biological interpretability and predictive accuracy?

To investigate these questions, we have developed ABMACT, a mechanistic modeling framework that reconstructs cellular dynamics of an evolving tumor-immune ecosystem, consisting of “virtual” immune cells and tumor cells defined by immunological knowledge and single cell molecular profiles obtained in experiments. We focused on NK cells in this first study and constructed an ABM with submodules of cellular functions and cell-cell interactions based on our biological knowledge of NK cells and experimental data.

## 2. Results

### 2.1 A cell-level mechanistic modeling framework for NK-ACT

ABMACT is a computer simulation framework for studying cell population dynamics and interactions in ACT using ABM (**Figure 1**). Cell agents, the building blocks of the ABMs, are determined based on domain knowledge of interacting cell populations (**Figure 1 Step1, Methods**)^2,12,13,15,45^. NK cells are *ex vivo* expanded to express activating receptors such as CD16, NKG2D, and activating Killer cell immunoglobulin-like receptors (KIRs) and are “licensed” to kill^12,45^. Cytotoxic killing is the primary mechanism determining therapeutic responses in ACT^15^, placing cytotoxic NK cells (*N*_*C*_) and tumor cells (B, such as B cell lymphoma) interactions in the center of modeling. However, activation and repeated killing can induce exhaustion, resulting in NK dysfunction and cancer immune evasion ^46^. While NK cells were believed to act short-term as part of innate immunity, multiple recent studies have highlighted their capacity to develop a “vigilant” phenotype – long living, dormant, and reactive to second pathogen stimulation^47–49^. To model the fate transitions of *N*_*C*_, we include exhausted NK cell agents (*N*_*E*_) and vigilant NK cell agents (*N*_*V*_). Cellular processes such as proliferation, exhaustion, death, antigen recognition, and migration have been characterized mathematically^24,41,50,51^. Drawing insights from previous studies, we further defined the specific mathematical rules and parameters in our NK cell agents using data obtained from *in vitro* autonomous growth and rechallenge assays (**Methods, Supplementary Material 1**). The increasing availability of single-cell molecular profiling data provides an unprecedented opportunity to model functional heterogeneity at cellular resolution. To achieve so, we quantified the effects of genes and pathways on cellular functions such as cytotoxicity. In this study, we used paired single-cell RNA-seq (scRNA-seq) and phenotype data from the xenograft lymphoma mouse models in Li et al.^23^ to select and estimate a subset of genetic features. Gene expression profiles are randomly assigned to the cell agents and translated to functional properties through the estimated effects. As a result, cell agents unbiasedly represent rich molecular profiles, modeling variations in individual cellular fates and collective populational dynamics (**Figure 1 Step 2, Methods**).

**Figure 1.**
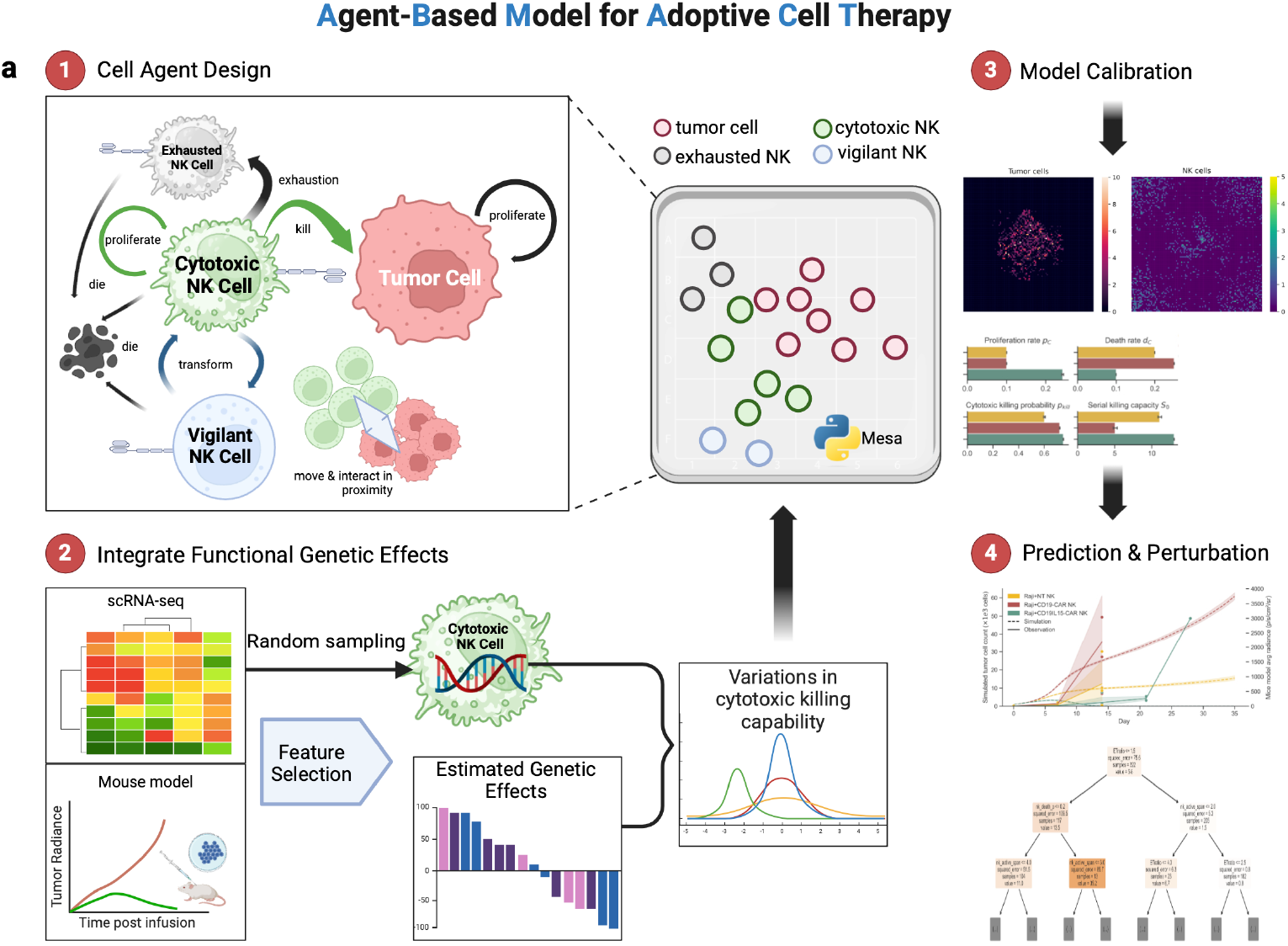
A mechanistic cell-level in silico modeling framework to elucidate cellular characteristics and cell population dynamics. a) Workflow diagram of Agent-Based Modeling for Adoptive Cell Therapy (ABMACT). Step 1: Cell agent design. Four cell types are included in ABMACT: cytotoxic NK cells, exhausted NK cells, Vigilant NK cells, and tumor cells. Step 2: Embed cell-level genetic effects. Functional effects of significant genes and pathways regulating NK cell cytotoxicity are derived from paired longitudinal scRNA-seq data and mice model experiment ^23^ (Methods 4.2). The coefficients, multiplied with randomly sample expression profiles in individual cytotoxic cell agents, contribute to variations in cytotoxic killing capability. Step 3: Cell agents interact in 2D simulated TME to model tumor-immune interactions over time. ABMs are calibrated on *in vivo* data. Step 4: Prediction and perturbation by modifying conditions in calibrated models.

We calibrated and evaluated the models using functional data obtained from various in vivo models such as the lymphoma mice model in Li et al. ^23^ and glioblastoma mice model in Shanley et al.^20^ (**Figure 1 Step 3, Methods**). By obtaining *in silico* replica of the *in vivo* systems, we can further perturb the models to examine the effects of various biological and therapeutic parameters and hypothetical conditions (**Figure 1 Step 4**).

### 2.2 ABMACT recapitulates differential tumor control in mouse models

We examined if ABMACT can model the course and the effects of *in vivo* NK cell therapy in mouse models (**Methods**). To further parametrize NK cell cytotoxicity from functional genetic effects, we applied feature selection to paired scRNA-seq and tumor radiance datasets from lymphoma mouse models in Li et al.^52^ using linear mixed effect regression (**Methods**). We identified significant gene and GO Biological Process (GOBP) pathway signatures that positively and negatively modulated NK cells’ anti-tumoral capacity, such as CD226, a known activating receptor for NK cell cytotoxicity^53^, and *PDCD1*, the Programmed Cell Death 1 gene, encodes the PD-1 protein that down-regulates NK cell anti-tumor activity^54^ (**Figure 2a, Supplementary Materials 2**). Cytotoxic NK cell agents were then initialized with randomly drawn scRNA expression to mimic the heterogenous NK cell population at the beginning of a treatment.

**Figure 2.**
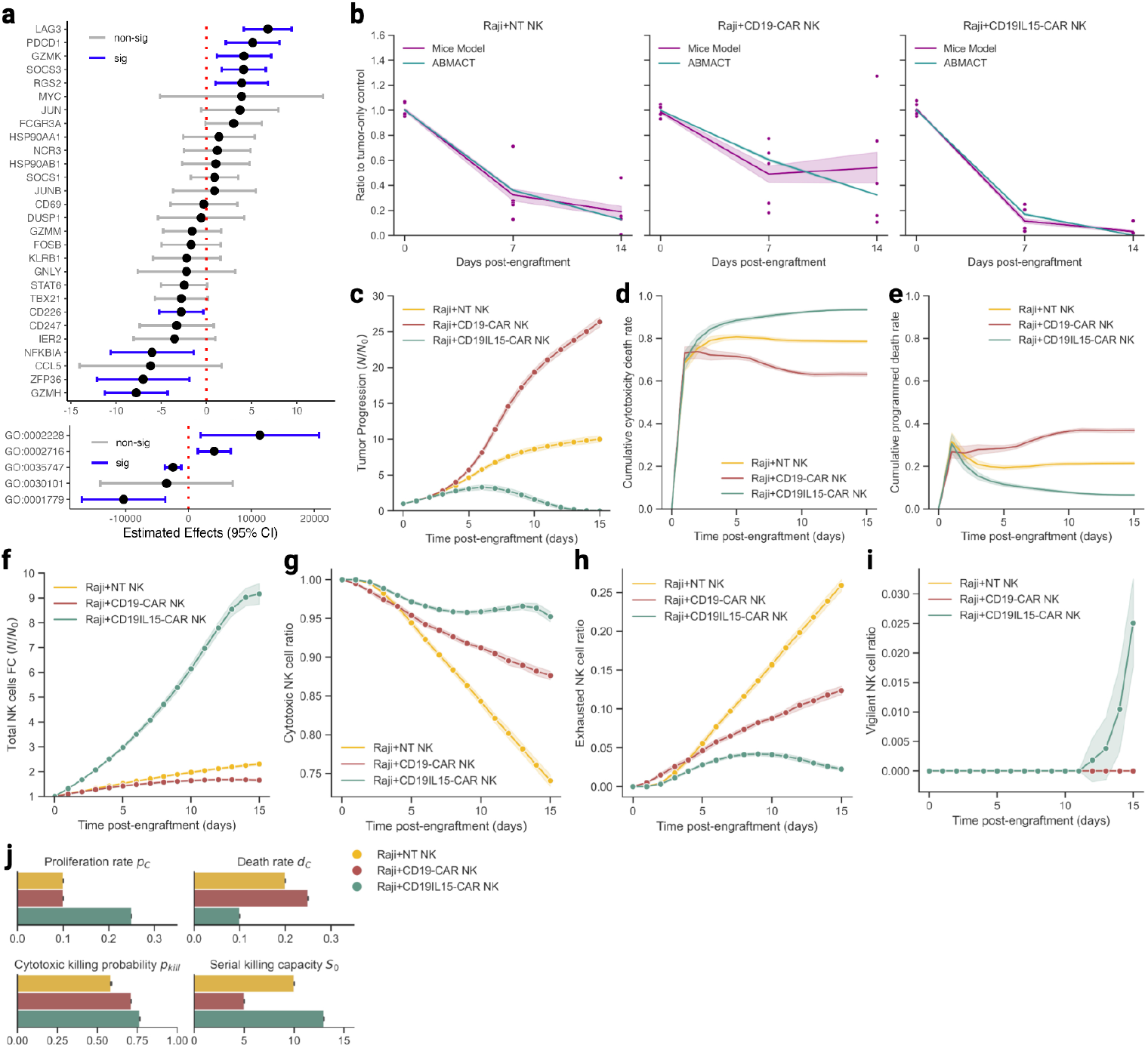
Modeling NK-tumor cell interactions in lymphoma mice models using ABMACT framework. (a) Estimated coefficients of significant genes and pathways regulating NK cell cytotoxicity (**Methods**). Confidence interval: 95%. (b) ABMACT simulations recapitulated normalized tumor dynamics in lymphoma mouse models. Comparison of simulated data with observed data using normalized tumor progression ratios between experimental groups and the tumor-only control group. (c) Tumor progression calculated by fold changes with respect to the initial tumor population at the start of the simulation. (d) The cumulative proportion of tumor cell death due to cytotoxic killing by NK cells. (e) The cumulative proportion of tumor cell death due to programmed death. (f) Total NK cell population fold changes with respect to the initial NK cell population. Normalized by NK cell count at day 0 of simulation when NK cells were added. Simulated ratios of (g) cytotoxic NK cells, (h) exhausted NK cells, and (i) vigilant NK cells with respect to the total NK cell population in the lymphoma model. (j) Fitted cytotoxic NK cell proliferation rate at baseline (t0 of simulation), death rate, cytotoxicity (probability of killing a tumor cell upon contact) at baseline, and serial killing capacity for three treatment groups. Results aggregated from the top ten fittings based on MSE. Interval bands of 2 s.e. were calculated using a bootstrap of 1000 iterations.

#### Lymphoma Mouse Model

We first investigated the therapeutic effects of engineered CAR-NK cells on a CD19+ lymphoblastoid cell line in immunodeficient mice in Li et al.^23^ (**Methods**). Best fitted parameter sets were found by iterations of global search and minimizing mean squared error (MSE). ABMACT achieved good fitting to the tumor volume data measured by bioluminescence imaging for all the three NK cell products: non-transduced NK (NT-NK), CD19 CAR-NK and CD19IL15 CAR-NK. Particularly, the CD19IL15 CAR-NK product showed more effective tumor control as compared to NT-NK and CD19 CAR-NK cells, with tumor clearance by day 14 post-engraftment (**Figure 2b**), recapitulating the *in vivo* experiments. With ABMACT’s ability to simulate continuous time courses, the differential tumor-killing capacity of the three NK cell products were depicted as continuous time trajectories, beyond the original time snaps (**Figure 2c**). ABMACT further delineated the cause of tumor cell death as NK cytotoxicity mediated death (**Figure 2d**) and programmed death (**Figure 2e**).

On the other hand, the NK cell population sizes appeared negatively associated with the tumor population size (Pearson’s R= **−**0E37, p<0.005, **Figure 2f)**. Notably, CD19IL15 CAR-NK cell population expanded at a faster rate, approached a plateau at day 15 after tumor clearance, while the other two products had slow, monotonic expansion (**Figure 2f)**. Cytotoxic NK cells, which constituted a large proportion of the total population throughout the 15-day simulation, had the fastest expansion in the CD19IL15CAR-NK cell group, contrasting with rapid decline in the CD19 CAR-NK cell and NT-NK cell groups (**Figure 2g**). The CD19IL15 CAR-NK cells also showed lower degree of exhaustion as compared with the other two groups (**Figure 2h**). However, lower exhaustion levels do not always coincide with more efficacious tumor control. Although CD19 CAR-NK cells appeared less exhausted than the NT-NK cells, tumors treated with the CD19 CAR-NK cells outgrew those treated with the NT-NK cells (**Figure 2c**). In the CD19IL15CAR-NK cell group, the vigilant phenotype emerged as tumor cells were cleared, indicating successful transition of surviving cytotoxic NK cells in the TME (**Figure 2i**).

We further attributed the NK cell dynamics to individual cellular properties. Based on parameters from the best-fitting results, we found that CD19IL15 CAR-NK cells had superior viability and killing capacity as compared to other two groups (**Figure 2j**). In addition to enhanced proliferation and lower death rates, higher cytotoxicity *p*_*kill*_ at the beginning of treatment, and higher serial killing capacity (SKC) *S*_0_, measured by the number of tumor cells one NK cell could kill before exhaustion, enabled CDIL15 CAR-NK cells to exert repeated tumor lysis at a high success rate. The systematic effects of the cellular properties resulted in stronger interactions between tumor cells and cytotoxic NK cells in the CD19IL15 CAR-NK cell treatment group than the other two treatment groups (**Figure S4a**). Interestingly, fitting results for NT-NK cells showed that the group had lower *d*_*C*_ and higher *S*_0_ than the CD19 CAR-NK cells, which potentially explained the observed relatively lower average tumor loads in NT-NKs than in CD19 CAR-NKs (**Figure 2b**). In lymphoma model, the optimal fitting results were found at an effector-to-target ratio (ETR) of 1:1, which was much lower than the set-up condition in the in vivo mice models (50:1). Under the ABMACT assumption of 100% cytotoxic NK cells in the initial NK cell population, such a difference in ETRs suggested that it was likely that only a fraction of NK cells infiltrated into tumor sites and exerted effector functions in the mice experiments.

From a modeling perspective, the inclusion of functional genetic effects in NK cell cytotoxicity is controlled by a genetic effect parameter *b*. Integrating scRNA-seq data by setting *b* > 0 improved the overall modeling accuracy (p-val=0.039), with the effects varied by group (**Figure S4d-e)**. In addition, embedding genetic effects enabled modeling variations in cytotoxic NK cells’ killing probabilities (**Figure S4c-d**).

#### Glioblastoma Mouse Model

To explore the potential of ABMACT in studying a broad spectrum of cancers, we further evaluated it in a glioblastoma (GBM) mouse model, which examines the therapeutic benefits of *ex vivo* expanded NK cells in kill GBM cell lines in immunodeficient mice (**Figure 3a**). The growth rate of GSC20 GBM tumor cells were estimated to be lower than Raji lymphoma (0.223/day vs 0.445/day), requiring a lower ETR (1:5 vs 3:1). Similar to the CD19IL15 CAR-NK cells in lymphoma model, cytokine-armed NK cells (IL-21 and IL-15 NK cells) showed more significant tumor control than NT-NK cells (**Figure 3b**). Tumor clearances were achieved in both IL-21 and IL-15 NK cells, with the highest proportions of cancer cell deaths induced by IL-21 NK cell cytotoxicity (**Figure 3c**). In the NT-NK cell group, tumor cells had sustained growth, with minimal reductions that were largely contributed by programmed deaths (**Figure 3d**). NK cells with cytokine-expressing vectors showed faster initial NK cell population expansion than the NT-NK cells (**Figure 3e**). However, the early cytotoxicity towards tumor cells in the IL-15 NK cell group also resulted in early drops in cytotoxic population (**Figure 3f**) and exhaustion (**Figure 3g**). The time of population shrinkage concorded with tumor clearance, which aligned with prior studies of NK cell dynamics^48,55,56^. Similar to CD19IL15 CAR-NK cells in lymphoma mouse model, a small proportion of vigilant NK cells emerged upon tumor clearance (**Figure 3h**). Despite having higher death rates than the IL-15 NK cells, the higher proliferation *p*_*C*_, cytotoxicity *p*_*kill*_, and SKC *S*_0_ of the IL-21 NK cells contributed to more rapid tumor control (**Figure 3i**).

**Figure 3.**
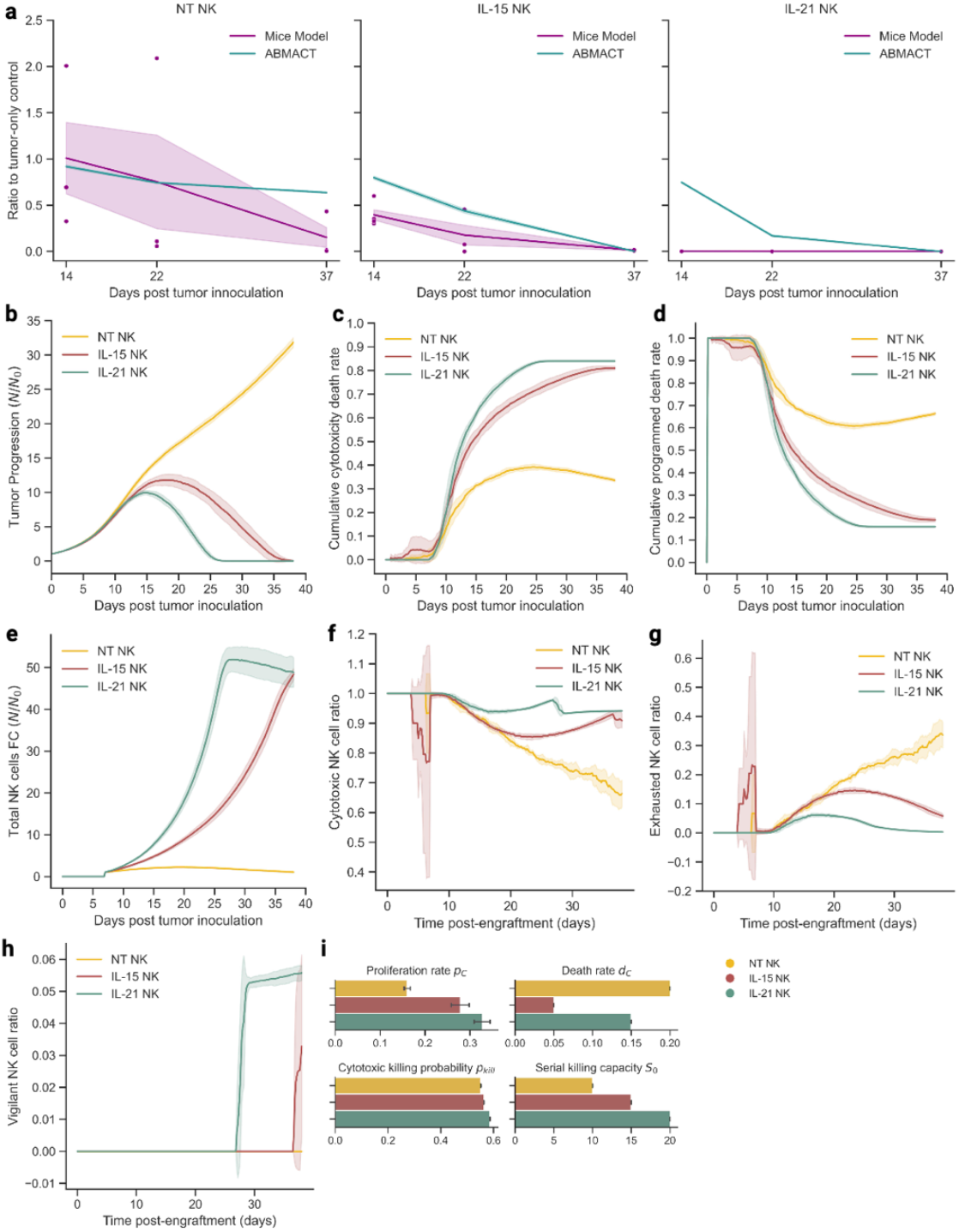
Modeling NK-tumor cell interactions in glioblastoma (GBM) mice models using ABMACT framework. (a) ABMACT simulations recapitulated normalized tumor dynamics in GBM mouse models. Comparison of simulated data with observed data using normalized tumor progression ratios between experimental groups and the tumor-only control group. (b) Tumor progression calculated by fold changes with respect to the initial tumor population at the start of the simulation. (c) The cumulative proportion of tumor cell death due to cytotoxic killing by NK cells. (d) The cumulative proportion of tumor cell death due to programmed death. (e) Total NK cell population fold changes with respect to the initial NK cell population. Normalized by NK cell count at day 7 of simulation when NK cells were added. Simulated ratios of (f) cytotoxic NK cells, (g) exhausted NK cells, and (h) vigilant NK cells with respect to the total NK cell population in the lymphoma model. (i) Fitted cytotoxic NK cell proliferation rate at baseline (t0 of simulation), death rate, cytotoxicity (probability of killing a tumor cell upon contact) at baseline, and serial killing capacity for three treatment groups. Results aggregated from the top ten fittings based on MSE. Interval bands of 2 s.e. were calculated using a bootstrap of 1000 iterations.

### 2.3 Applications of ABMACT

#### 2.3.1 Augmenting experimental observations by predicting treatment courses

*In vivo* systems are inherently limited by the frequency and resolution of measurements, often capturing only snapshots of dynamic treatment courses. One key advantage of *in silico* models is their ability to augment and complement existing experimental results, enhancing scale and granularity. ABMACT builds upon this strength by simulating designated durations and inferring subpopulation dynamics that maximally explain laboratory observations. For example, using models for CD19IL15 CAR-NKs, CD19 CAR-NKs, and NT-NKs calibrated on the lymphoma mouse model dataset in Li et al.^23^, we projected tumor progression and NK cell population dynamics beyond the endpoint of experimental observation (post mice sacrifice), extending the course to 35 days post-treatment (**Figure 4a-b**).

**Figure 4.**
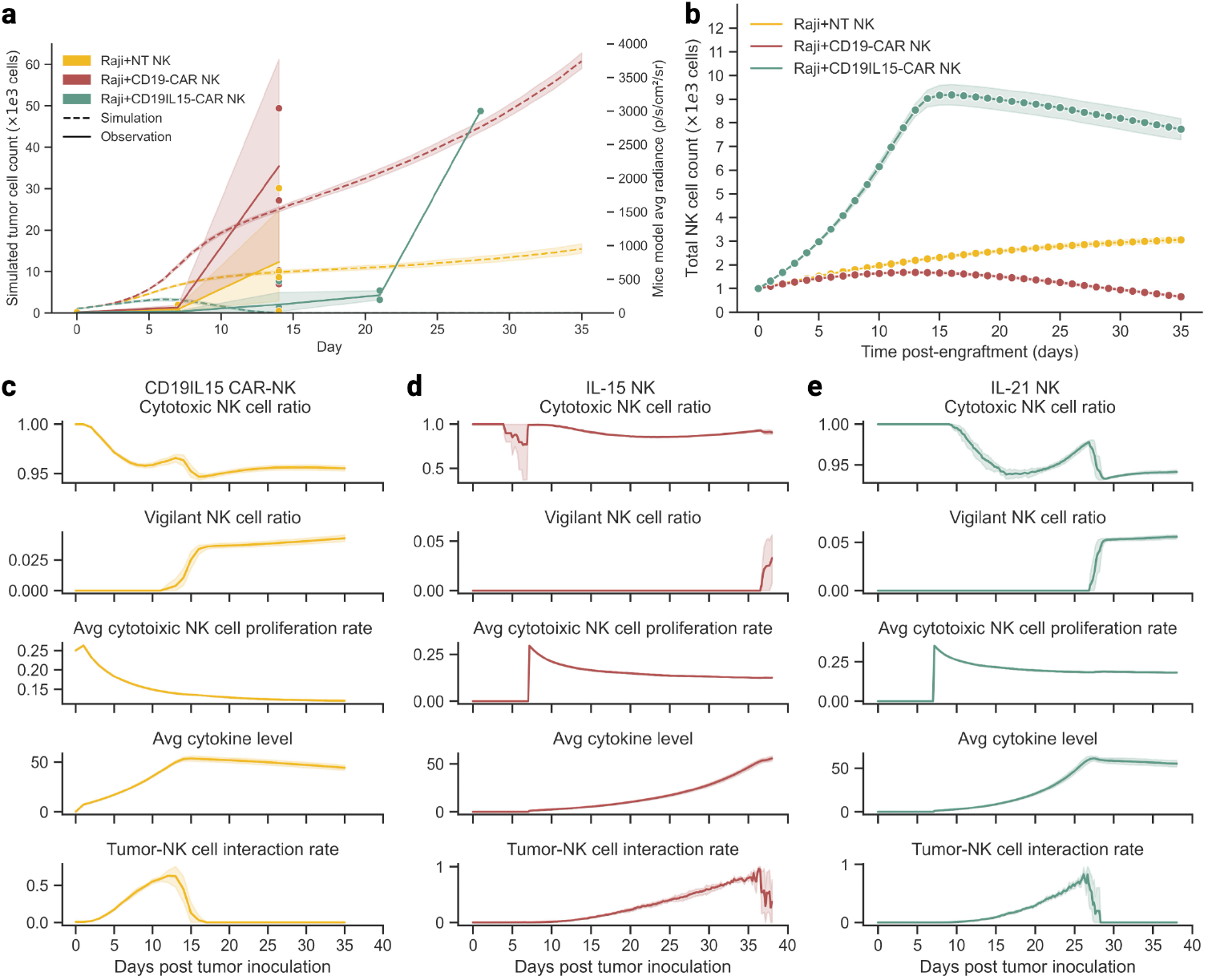
Plotting the course of cellular treatment using ABMACT. Prediction of tumor and NK population dynamics: (a) Simulated and observed tumor progression in the lymphoma mouse models. (b) Simulated total NK cell population trends in NT-NKs, CD19-CARNKs, and CD19IL15-CARNKs treatment groups in the lymphoma mouse model. Tumor/NK survival calculated by fold changes with respect to the initial tumor population at the start of the simulation. Deconvolution of phenotypes and kinetics: Cytotoxic NK cell ratio, vigilant NK cell ratio, average cytotoxic NK cell proliferation rate, average cytokine level, and tumor-NK cell interaction rate of (c) CD19IL15 CAR-NK cells in the lymphoma model, (d) IL-15 NK cells in the GBM model, and (e) IL-21 NK cells in the GBM model. NK cell subtype ratio calculated with respect to the total NK cell population at each timepoint. Tumor-NK interaction rate calculated by the ratio of tumor cells with co-locating cytotoxic NK cells with respect to the total tumor cell population.

Our model projected that the CD19IL15 CAR-NK treatment would successfully eliminate tumor cells without relapse before day 13. In contrast, tumor populations in NT-NK and CD19 CAR-NK cell groups would grow 15 to 60 folds from the initial 1000 simulated cells, respectively. Interestingly, the lower tumor growth observed in the NT-NK group compared to CD19 CAR-NK group and eventual tumor elimination in NT-NKs diverged from the expectation that NT-NKs would perform worse than CAR-engineered NK cells. However, this finding aligns with observations from the original lymphoma mouse model experiment, where the NT-NK group exhibited lower tumor radiance than the Raji-only group on day 14 post-engraftment (equivalent to day 21 post-infusion) (**Figure S4a**). These seemingly counterintuitive results underscore the need for further investigation into the functional properties and mechanisms underlying the performance of the NT-NK and the CD19 CAR-NK treatments.

By deconvoluting NK cell populations in the lymphoma and the GBM mouse models, we identified distinct kinetics of engineered NK cells. Notably, cytokine-expressing NK cells, including CD19IL15 CAR-NKs (**Figure 4c**), IL-15 NKs (**Figure 4d**), and IL-21 NKs (**Figure 4e**) exhibited a small peak in cytotoxic NK cell ratios, corresponding to the transition of a subset of cytotoxic NK cells into the vigilant phenotype upon tumor clearance. In contrast, this pattern was not observed in NT-NKs and CD19 CAR-NKs (**Figure S5a-c**). This phenomenon was likely driven by the co-stimulation of endogenous cytokines and residual tumor presence, indicated by cytokine levels and declining tumor-NK cell interaction rate. The cytotoxic NK cell population sustained proliferation while were less prone to exhaustion at the phenotype shift. While the NK cell phenotypes defined in ABMACT are hypothetical and informed by existing studies, the observed NK cell subpopulation trajectories may provide insight into treatment efficacy and warrant further experimental validation.

#### 2.3.2 Discovering key drivers of tumor control through *in silico* perturbation experiments

While numerous CAR engineering strategies aim to enhance NK cells, exhaustive testing of potential designs require significant amount of time and resources. To prioritize the most critical factors influencing tumor control in NK-ACT, we conducted feature importance sensitivity analysis (**Methods**) using *in silico* perturbations in NK cell properties and dosages. The analysis revealed that the effector-to-target ratio (ETR) was the most important feature for accumulated tumor growth, followed by NK cell serial killing capacity (SKC) *S*_0_, death rate *d*_*C*_, baseline proliferation rate *p*_*C*_ and its decay rate 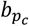, and baseline cytotoxicity *μ*_*C*_ (**Figure 5a, Figure S5d**). Increasing ETR from 1:1 to 2:1 drastically shortened the time to tumor clearance, though further increases yielded diminishing returns (**Figure 5b**), aligning with findings previously reported in a CAR-T dosing review study^57^. On the contrary, enhancing SKC and proliferation rates showed a continuous trend of accelerated tumor clearance (**Figure 5c-d**). Parameters contributing to NK cell killing, including *S*_0_, *μ*_*C*_, CAR effect exponent *γ*, and cytotoxicity genetic effect coefficient *b* had a combined feature importance 61% higher than the combined importance of viability parameters 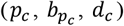, suggesting that enhancing *N*K cell killing capacities such as SKC, cytotoxicity, and specific recognition may be more effective than improving their viability in the system. We also found that higher NK cell mobility, parametrized by the probability to move at every step of simulation (*m*_*N*_) further reduced the time to tumor clearance (**Figure S5e**).

**Figure 5.**
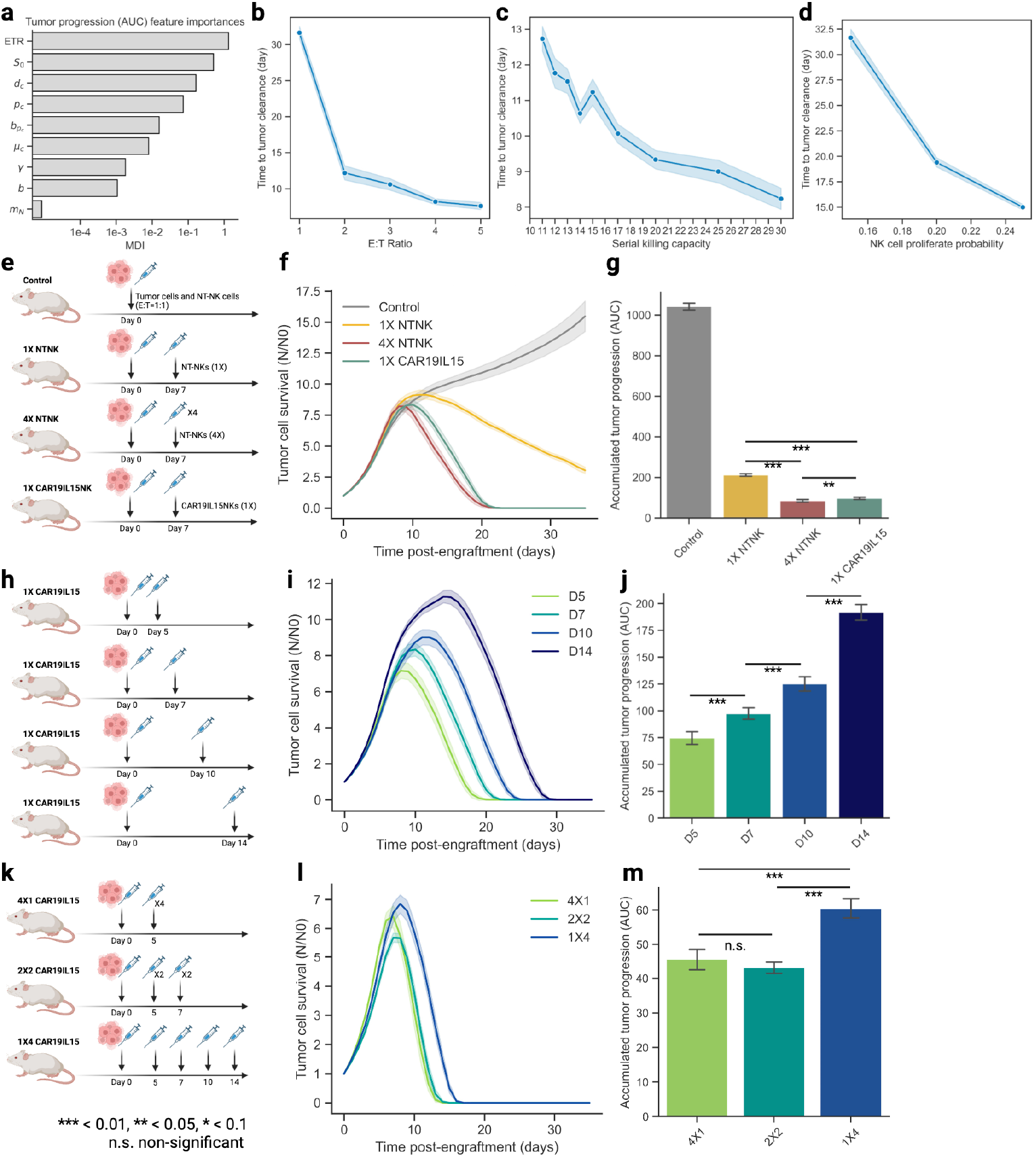
Predicting optimal treatment regimens using ABMACT. Sensitivity analysis: (a) Random Forest regression feature importance of model parameters on tumor progression area under the curve (AUC) measured by mean decrease in impurity (MDI) (**Methods**). Time to tumor clearance by (b) effector-to-target ratio, (c) serial killing capacity, and (d) NK cell baseline proliferation rate. Other parameters and simulation conditions were kept constant. Simulating NK-ACT follow-up treatment: (e) Schema of virtual NK-ACT treatments. Control: NT-NKs were administered at day 0 at an E:T ratio 1:1. 1X NTNK: One follow-up dose (1X) of NT-NK cells was administered on day 7. 4X NTNK: Four follow-up doses (4X) of NT-NK cells were administered on day 7. 1X CD19IL15 CAR-NK: One follow-up dose (1X) of CD19IL15 CAR-NK cells administered on day 7. (f) Simulated tumor progression and (g) accumulative tumor growth (Area under the curve, AUC) of control, 1X NTNK, 4X NTNK, and 1X CD19IL15 CARNK cell treatments. (h) 1X CD19IL15 CAR-NK: One follow-up dose (1X) of CD19IL15 CAR-NK cells was administered on day 5, 7, 10, and 14. (i) Simulated tumor progression and (j) accumulative tumor growth (AUC) of the four treatment groups. (k) 4×1 CD19IL15 CAR-NK: Four follow-up doses (4X) of CD19IL15 CAR-NK cells administered on day 5. 2×2 CD19IL15 CAR-NK: Two follow-up doses (2X) of CD19IL15 CAR-NK cells administered on day 5 and 7 each. 1×4 CD19IL15 CAR-NK: One follow-up dose (1X) of CD19IL15 CAR-NK cells was administered on day 5, 7, 10, 14 each. (l) Simulated tumor progression and (m) accumulative tumor growth (AUC) of the three treatment groups. 10 simulations for each experiment. Interval bands of 2 s.e. were calculated using a bootstrap of 1000 iterations.

#### 2.3.3 Assisting treatment decision-making using ABMACT simulation

ACT dose responses often deviate from linear relationships, where excessive dosages may fail to significantly improve treatment efficacy while increasing the risk of toxicity ^57^. Designing optimal ACT treatment regimens is a delicate task, with limited room for repeated experimental testing. ABMACT provides a valuable platform for exploring treatment options and informing clinical decision-making through in-silico simulations. In the lymphoma mouse model simulation, a single dose (1:1 ETR) of NT-NKs at the ETR of 1:1 led to tumor outgrowth (**Figure 2c**). To investigate effective tumor control strategies, we evaluated follow-up treatments with varying dosages, frequencies, timing, and NK cell products using ABMACT simulations (**Figure 5e**). We found that to effectively control tumor outgrowth, it required a higher dose of the same NK cell products (4X) or more effective NK cells such as CD19IL15 CAR-NK cells (1X) (**Figure 5f**). The overall tumor growth was significantly lower in the 4X NT-NK cell group (p-adj < 0.001) and 1X CD19IL15 NK group (p-adj < 0.001) (**Figure 5g**).

Next, we investigated whether administrating the follow-up treatment at different times would result in differential tumor control. One dose (1X) of CD19IL15 CAR-NK cells was administered on day 5, 7, 10, and 14 following the initial treatment, respectively (**Figure 5h**). Earlier intervention contributed to more efficient tumor control (**Figure 5i**) and significantly smaller cumulative tumor burden (**Figure 5j**). Treating refractory tumors with higher doses of more potent ACT products can lead to a more rapid response, and dose fractionation have been used in CAR-T cell therapies and recombinant radiotherapy with CAR-NK cell therapy to mitigate the risks of adverse events associated with high dosages^58,59^. However, it is unclear whether high dosing in NK-ACT is associated with risks^60^, and fractioned dosing in NK-ACT alone has not been extensively explored. To examine whether dose fractionation in NK-ACT can effectively control tumors, we simulated follow-up treatment with a total of four doses of CD19IL15 CAR-NK cells administered in one treatment, two treatments, and four treatments (**Figure 5k**). No significant difference in cumulative tumor growth was shown when administrating four doses in one treatment versus splitting to two treatments, but these two treatment strategies resulted in earlier tumor clearance and smaller cumulative tumor burden compared to splitting administrating the four doses over a course of four treatment (**Figure 5l-m**). For aggressive cancer types such as lymphoma, tumor cells escaping NK cell surveillance gain a head-start and undergo exponential growth^61^. When initial treatment fails, higher doses or more potent NK cell products are often required to regain control. The timing of treatment influences response efficacy, while fractionated dosing under calibrated dosage and timing can potentially provide the same treatment benefit as a single higher dose. ABMACT offers a predictive framework for pretesting follow-up treatment strategies, reducing the reliance on extensive laboratory experimentation.

## 3. Discussions

Recent progress in natural killer (NK) cell adoptive cell therapies (ACT) for cancer treatment have sparked significant interest, yet the underlying cellular kinetics, cell-cell interactions, and molecular mechanisms remain incompletely understood. NK cell engineering involves addressing complexities in intrinsic cellular properties, TME heterogeneities, and needs for extensive experimental validations. Although *in vitro* and *in vivo* models are indispensable, they are limited in variety, scalability and resolution. Mechanism-based computational models, particularly ABMs, can provide an efficient and ethical alternative^62^.

We developed ABMACT, an agent-based modeling framework for NK-ACT, incorporating biological rules derived from experimental data, literature, and single-cell molecular profiling. By studying tumor-immune dynamics through simulations, we found better-performing NK cell products possessing superior cellular properties including higher viability, cytotoxicity, and serial killing capacity, i.e. CD19IL15 CAR-NKs in treating lymphoma and IL21 NKs in treating GBM. Similar cellular properties of NT-NKs were observed in both mice experiments, while NK cell products with varying engineering strategies showed distinct characteristics. Administration strategy and cancer type also lead to differences in tumor dynamics. Simulating from engraftment of NK cells to tumor locations, the lymphoma model exhibited differences in tumor control in the early phase of simulation. Such differences were manifested later in the GBM model, where NK cells were administrated seven-day post-tumor inoculation rather than through blood vessel diffusion. The required ETR was lower in the GBM model than the lymphoma model, given the less aggressive cell line used in the PDX model. Overall, cell behaviors, cancer conditions, and therapy administration strategies were virtually mimicked in the ABM, facilitating detailed investigation.

Compared with other modeling techniques such as ODE, ABMACT provided enhanced accuracy and details. The overall accuracy measured by mean squared error of the ABMs appeared comparable to those obtained from ODE models and outperformed in two out of three cases (**Figure S4b-c, Supplementary Materials 5, Table S5**). In addition to accuracy, an important aspect of computational modeling is interpretability^62^. ABM provided advantages in recapitulating and decomposing experimental observations across various contexts, including NK cell autonomous growth, rechallenge experiments, and lymphoma and glioblastoma mice models. Our framework allows quantitative assessments of NK cell cytotoxicity, exhaustion, and long-term viability, generating insights and hypotheses for ACT treatment efficacy.

Another practical advantage of ABMACT is its implementation using the MESA Python framework^63^, ensuring portability, accessibility, and compatibility with existing computational tools. Reproducibility of the results can be easily warranted. As the field of ABMs expands, institutions and individuals can conveniently acquire cell agent templates and modify their functions and properties. Surrogate models, such as Gaussian process regressors, can further enhance simulation speed when rapid predictions are prioritized over granularity^62^. These features make ABMACT an adaptable and user-friendly platform for researchers, facilitating hypothesis generation, pre-testing, and iterative improvements to experimental design.

A major challenge in cancer treatment is the dynamics in cellular physiology throughout treatment courses, complicating the prediction of treatment outcomes and clinical decision-making^24^. A key feature of ABMACT is its ability to simulate diverse NK cell engineering designs and treatment regimens. Aggregating data from over hundreds of simulation scenarios, we found that in addition to sufficient immune cell engraftment to the tumor site, NK cell serial killing capacity, cytotoxicity, and proliferation capacity emerged are critical determinants of effective tumor control. We demonstrated that in addition to the intrinsic efficacy of cell products, dosages, and timing also influence treatment prospects. *In silico* perturbations enable virtual testing of treatment plans, reducing reliance on resource-intensive and ethically challenging animal experiments. Moreover, integrating statistical modeling enhances ABMACT’s robustness by accommodating variability beyond what can be achieved using cell-line or animal models alone. With ABMACT, simulations can be run to obtain individualized, continuous prediction of tumor regression/progression and treatment outcomes, which is highly useful for risk assessment and treatment personalization.

Building on these capabilities, ABMACT offers the potential to complement preclinical and clinical applications, streamlining therapeutic development and optimizing treatment strategies. ACT, particularly those involving engineered NK cells, show promise in targeting a range of malignancies. ABMACT’s ability to simulate dosage strategies, administration timing, and engineered cell designs provides a platform for preclinical hypothesis generation. These simulations can help refine critical parameters, such as effector-to-target ratios or serial killing capacities, and inform clinical trial design, reducing reliance on labor-intensive in vivo experiments,

Challenges in ACT, such as tumor heterogeneity, immune-evasive clones, and patient-specific variability, require careful consideration. ABMACT can incorporate molecular profiling and patient-derived data to simulate these factors and propose strategies for personalized treatment. For instance, patient-specific cytokine profiles could enhance predictions of NK cell dynamics and treatment outcomes. However, it is important to note that computational models like ABMACT are tools to complement experimental and clinical research, not replacements, Predictions must undergo validation, and their role is to guide, not dictate, clinical decisions. By integrating computational insights with experimental findings, ABMACT has the potential to enhance ACT development while maintaining a realistic perspective on its applicability.

Parametrizing ABMs depends on data availability and requires balancing computational trackability, and modeling accuracy. Despite its strengths, the current version of ABMACT has several limitations. Firstly, while addressing cellular and molecular dynamics of NK-tumor interactions, it does not fully account for the immune system and host-level biology as a single-compartment model. ABMACT captures the major cell types but has not included the remaining immune cells such as monocytes, neutrophils, and macrophages in NSG mice. Deviations in simulations of CD19CAR-NKs in mice model from the observed data after day 14 suggested residual tumors due to inadequate NK cell infiltration to tumor engraftments or incomplete elimination. That is, tumor cells were likely to persist in multiple lymph nodes where they evaded immune surveillance. In the same mouse model, cytokines expressed by CAR-NKs also led to toxicity and deaths of experiment animals. Future work will be needed to address these issues. Secondly, despite improvement to model accuracy and the ability to capture variations in NK cell cytotoxicity driven by gene and pathway markers, the small sample size of paired longitudinal scRNA-seq and tumor radiance data from mouse models may reduce effect size and lead to the omission of key markers. ABMACT can benefit from aggregating datasets from multiple studies and establishing genetic markers that characterize various NK cell functionalities beyond cytotoxicity with higher statistical power. Thirdly, our current model does not consider treatment toxicity, as NK-ACTs have generally been associated with little adverse effects in previous clinical trials^1,5^.

The future work of ABMACT can expand current modeling of cellular and molecular dynamics and extend beyond its initial application to NK-ACT. NK cell activities can be decomposed to viability, cytotoxicity, exhaustion, and memory-like adaptation, while NK cell metabolism also contribute to its effector function^23^. Data such as OXPHOS, glucose consumption, and TME pH can be incorporated in ABMACT to characterize metabolic competition between NK cells and tumor cells. The modular design supports expanding cell agent and TME specifications. Cell types such as macrophages, CD4+ and CD8+ T cells, and dendritic cells are known to play crucial roles in anti-tumor immunity. They can be modeled by modifying and extending cell agent design in the current ABMACT. The modular structure of ABMACT allows modification of tumor kinetics, modes of immune cell infiltration, TME cytokine pharmacokinetics, and various aspects of cell properties, enabling flexible extension to other tumor types, physiological conditions, and multi-compartmental biology. Spatial transcriptomics and multiplexed imaging data can also be integrated in ABM to provide positional priors for cells as previously demonstrated ^61^. Additionally, coupling ABMs with ODEs or partial differential equations (PDE) could model signaling cascades and gene regulatory networks^64^. Such hybrid models can reduce computational costs while maintaining biological relevance, capturing dynamics at both cellular and molecular levels, and extending ABMACT’s applicability to broader contexts of immunotherapies.

In conclusion, ABMACT represents a significant advancement in the computational modeling of NK cell therapies. By integrating experimental data and single-cell transcriptomics, it provides mechanistic insights into tumor-immune interactions and highlights critical cellular properties underpinning treatment efficacy. As directed by the FDA Modernization Act 2.0^65^, new alternative methods are sought to reduce the necessity of animal testing. The knowledge-informed, ruled-based parametrization in ABMs allows investigating new therapies when limited data are available. Future studies should focus on synchronized planning of laboratory experiments and computational modeling to facilitate hypothesis generation and accelerate new drug development. In clinical investigations, ABMs can be useful in incorporating patient data to generate personalized treatment course prediction and potentially to optimize treatment regimen. This can potentially accelerate therapeutic development and assessment and eventually lead to improved clinical outcomes. As the first agent-based model dedicated to engineered NK cell therapies to date, the step-by-step workflow demonstrated in this study can be further extended with other data modalities, cell types, and cancer pathologies, paving the way for the development of more robust and personalized computational tools to guide the design and optimization of adoptive cell therapies.

## 4. Methods

### 4.1 In vitro and in vivo experiment data

#### 4.1.1 In vitro cell autonomous growth

Cell counts of NT (n=3) and CD19IL15 CAR-NK (n=3) from day 0 to day 42 were obtained from the cord blood NK cell autonomous growth experiments in Liu et al.^19^. Cells were cultured in vitro without tumor or additional cytokine stimulation.

#### 4.1.2 In vitro tumor rechallenge assay

##### Raji lymphoma

Mean tumor population dynamics measured by tumor cell index were obtained from tumor rechallenge assays with cord blood CAR19/IL15 NK cells from optimal cords (Opt-Cs) and suboptimal cords (Sub-Cs) (n=4 each) in Marin et al.^5^. The tumor cell index was measured by the intensity of mCherry fluorochrome, representing the counts of tumor cells. NK cells were challenged against mCherry transduced Raji lymphoma tumor cells at an effector-to-target ratio (ETR) of 5:1. Tumor cells (100,000 cells) were added every two to three days.

##### Glioblastoma

Mean tumor population dynamics measured by tumor cell index were obtained from GSC20 tumor rechallenge assays with IL-21 and IL-15 NK cells (n=3 donors each) in Shanley et al.^20^.NK cells were challenged against mCherry transduced GSC20 glioblastoma cells at an E:T ratio of 1:1. Tumor cells were added every two to three days.

#### 4.1.3 In vitro dose-response assay

Mean cytotoxicity profiles of NT, CAR19, CAR19/IL15 NK cells were obtained from the ^51^Cr-release dose-response assay of NK cell products (n=3 donors) against Raji targets in Li et al.^23^. Cytotoxicity was measured as the percentage of specific lysis of tumor cells relative to targets.

#### 4.1.4 Xenograft lymphoma mice model

##### Lymphoma

Tumor growth data and scRNA-seq data of NK cells and tumor cells were obtained from the xenograft mice model of Raji lymphoma treated with NK cells in the Figure 2C of Li et al.^23^. The scRNA-seq data are publicly available in Gene Expression Omnibus (GEO) repository at accession number GSE190976. Tumor loads were quantified as average tumor radiance in p/s/cm^2^/sr, which were assumed to be proportional to size of the tumor population (**Figure S3a**). Normalized tumor progression was calculated by dividing tumor radiances of experiment groups by the mean tumor radiances in the tumor-only control group (**Figure S3b**). NK cells were transfected with retroviral vectors encoding iC9.CAR19.CD28-zeta-2A-IL-15 (CAR19IL15), CAR19.CD28-zeta (CAR19), and IL-15, with non-transduced (NT) NK cells serving as control (n=5 per group). The xenograft NOD/SCID IL-2Rγ null mice were infused with FFLuc-labeled NK-resistant Raji lymphoma cells (2 × 10^5^ per mouse) on day 0. NK cells were harvested from each group pre-infusion and on days 7, 14, 21, 28 and underwent scRNA sequencing. Tumor cells were harvested from each group on days 7, 14, 21, 28 and underwent scRNA sequencing.

##### Glioblastoma

Tumor growth data and scRNA-seq data of NK cells were obtained from the xenograft mice model of GSC20 glioblastoma treated with NK cells in Figure 3A of Shanley et al.^20^. The processed scRNA-seq data are publicly available in Gene Expression Omnibus (GEO) repository at accession number GSE227098. Tumor loads were quantified as average tumor radiance in p/s/cm^2^/sr (**Figure S3c**). Normalized tumor progression was calculated by dividing tumor radiances of experiment groups by the mean tumor radiances in the tumor-only control group (**Figure S3d**). The NOD/SCID IL-2R-null (NSG) human xenograft mice were intracranially injected 0E5 × 10^6^ patient-derived FFluc-labeled GSC20 tumor cells on day 0 and treated intratumorally with NK cells (n = 3 to 5 per group) at an E:T ratio of 1:5 at day 7.

### 4.2 Cell agent design

We encoded three NK cell phenotypes: cytotoxic NK cells (N_C_), exhausted NK cells (N_E_), vigilant NK cells (N_V_), along with tumor cells (B, such as B cell lymphoma) in the agent-based model (ABM) using the Python Mesa framework^63^ (**Figure 1, Figure S1**). The tumor microenvironment was established in a 2D Moore neighborhood discrete lattice grid, which is commonly used in ABM, given its simplicity and efficiency in modeling^44^. Cell Agents act autonomously by programmed rules and commit to an action when the probability, sampled from a uniform distribution *U* (0,1), passes a predefined threshold e.g. proliferation rate. For example, Raji lymphoma tumor cells were estimated to have a proliferation rate of 0.455/day. When at a completion of a cell cycle, a tumor cell agent samples a random number from *U* (0,1) and compare it with 0.455. If the random number exceeds the threshold, the cell will divide and generate a daughter cell. Cellular properties are encoded as attributes of agents and are inherited by daughter cells from mother cells. The TME was initiated with tumor cells in the center and NK cells in the periphery. We assumed that cell agents followed Brownian motions and modeled cell motility with random walks, while NK cell agents traveled in the direction of the highest tumor concentration due to the chemokine gradient when tumors are present. The specifics of cell agent design are entailed below.

#### 4.2.1 Cytotoxic NK cells (*N*_*C*_)

A cytotoxic NK cell interacts with a target in three stages: migration, conjugation, and attachment^66^. NK cells can both move freely and migrate towards chemokine or proinflammatory protein gradients^67^. When reaching and recognizing the target, the conjugation phase starts. The NK cell forms an immunological synapse and reorganizes actin cytoskeletons^68,69^. The microtubule organizing center (MTOC) and secretory lysosomes are polarized towards the immunological synapse, followed by lysosome docking and finally the release of cytotoxic molecules into the target cell^68,69^. On ending the conjugation phase, the NK cell begins to dissociate from the target cell irrespective of successful killing, resuming free migration or initiating conjugations with other targets^66^. A cytotoxic NK cell (*N*_*C*_) is responsible for killing tumor cells. When a *N*_*C*_ cell agent encounters a tumor cell agent, *N*_*C*_ has a probability of *p*_*kill*_ to successfully kill the tumor cell, depending on various factors contributing to its cytotoxicity and the tumor cell’s ability to evade immune surveillance (*p*_*evade*_). In the process of killing, *N*_*C*_ reduces its serial killing capacity (*s*), resulting in exhaustion and transformation to the exhausted phenotype (*N*_*E*_) when *s* reaches 0. As compared to the other two NK cell phenotypes, *N*_*C*_ cell agents are able to efficiently expand and have the potential to transform to a vigilant phenotype (*N*_*V*_). In Vanherberghen *et al*.^66^, the mean total conjugation and attachment time was measured to be 193 minutes and 235 minutes for lytic and non-lytic events, respectively. Therefore, we considered a four-hour step length (Δ*T* = 4*hr*) for the ABM to approximate the time for cytotoxic NK cells moving toward tumor cells and exert killing.

##### 4.2.1.1 Cytotoxic killing activity

We constructed the probability of killing a tumor cell upon contact *p*_kill_ with reference to the cytotoxicity function in ^41^. *p*_*kill*_ is a function of baseline cytotoxicity c_NK_, gene effects G_NK_, CAR engineering effect *γ*, and the tumor cell’s probability to evade recognition p_evade_:

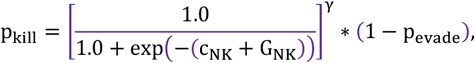

where 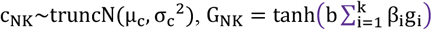, and b is the overall NK cell cytotoxicity genetic effect coefficient to ensure reasonable scaling. The initial value of NK cell baseline cytotoxicity *μ*_c;_ was estimated based on the dose-response data from the ^51^Cr-release assay of NK cell products against Raji targets in Retzlaff et al.^23^ and refined using global search (**Supplementary Material 1**). The RNA expressions of significant genes and pathways associated with NK cell cytotoxicity **g** and respective coefficients **β** were obtained from the cross-lagged LME model *M*_*NK*_.

##### 4.2.1.2 Characterize NK cell proliferation kinetics

NK cells require extrinsic stimulations to expand and persist. Studies have shown that the presence of cytokines such as IL-15 and tumor antigens enhances NK cell expansion^5,19,23^. Without exogenous stimulation, NK cell expansion could not be sustained, and the population start to wane in one to two weeks^70^. In the cell autonomous growth experiment in Liu et al.^19^, IL-15-expressing CAR-NK cells sustained higher population than NT NKs, although the effect of IL-15 stimulation on NK cell proliferation and survival gradually reduced due to system clearance^19^.

Therefore, we hypothesized that the endogenous cytokine expression such as IL-15 in CAR-NK cells is crucial for population expansion in addition to tumor antigen stimulation. Using an exponential form and Hill’s equation to describe the nonlinear dependencies between cytokine concentration and cell proliferative property^71^, we proposed a cytokine-dependent model (CM) for computing the proliferation rate of IL-15-expressing NK cells as follows.

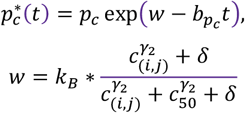

Initial ranges and default values of hyperparameters were determined using linear programming of biological constraints. *p*_*C*_ is baseline cytotoxic NK cell proliferation rate, *c*_(*i,j*)_ is the dimensionless cytokine concentration level in the neighboring region (*i, j*), *c*_50_ is the half-maximum cytokine level. *γ*_2_ is the Hill’s equation exponent, 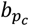 is the natural decay rate parameter of proliferation rate, *t* is cell age, and *δ* is a small constant for keeping *w* nonzero 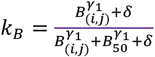 is the tumor stimulation effect considering tumor count *B*_(*i,j*)_ in the neighborhood (*i, j*), half-maximum tumor load *B*_50_, and a Hill equation exponent *γ*_1_. *k*_*B*_ is a constant in the context of NK cell autonomous growth as tumor cells are absent and kept to the default value. We considered a half-life of 2.5hrs for IL-15^72^. At every step of the ABM, the real-time cytotoxic NK cell proliferation rate 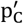 was sampled from a 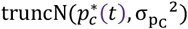 distribution.

CM is compared to a vanilla model (VM) of *w* = 0 and a *b* invariant across groups. The first 70% of data were used for fitting and the remaining for testing. Parameters *c, γ*_2_, *b* were globally searched to minimize mean squared errors (MSE) between simulated and observed population fold change with respect to the initial cell counts. The characterization of NK cell proliferation model is performed on the autonomous growth kinetics of cord blood (CB) CAR-NK and NT-NK cells using measured cell counts data from ^19^. Results are provided in **Supplementary Material 1**.

#### 4.2.2 Exhausted NK cell (*N*_*E*_) and characterize NK cell exhaustion process

In adoptive cell therapy, NK cell serial killing capacity (SKC) can be modulated by its CAR engineering^23^, source quality^5^, gene editing^22^, and cytokine stimulation^73^. Once a cytotoxic NK cell exhausts, it will no longer be able to kill tumor cells but can remain in the system in the presence of tumor antigen. We focused on aspects of NK cell exhaustion due to loss of ability to secret cytolytic granules and impaired cytotoxicity due to dysregulated inhibitory signals. The former was quantified as the serial killing capacity *s*, and the latter was modeled by a link function between the probability of killing a tumor cell upon contact, *p*_*kill*_ and RNA expression of genes found to be significantly associated with NK cells’ tumor control ability based on the LME modeling. Bhat et al.^73^ demonstrated that one NK cell was able to kill four to six tumor cells in 16 hours. However, the experiment was limited in duration to thoroughly measure the maximum SKC. To date, the exhausted NK cell phenotype can only be determined functionally using NK cell rechallenge assay. To fill the gap of mathematical models of NK cell exhaustion, we hypothesized that NK cell exhaustion could be deconvoluted as the reduction of serial killing capacity and impaired killing capability. We encoded an *NK cell initial SKC* parameter S_0_, which reduced as NK cell killed targets (event denoted as *I*_*kill*_). The following Exhaustion Models (EM) are proposed.

Exhaustion Model 1 (EM1):

NK cell exhaustion can be primarily described by the linear reduction in SKC. A cytotoxic NK cell (*N*_*C*_) transformed to an exhausted NK cell (N_E_) when its current SKC reached zero.

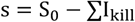

Exhaustion Model 2 (EM2):

In addition to the linear reduction in SKC, interactions with tumor cells increase the inhibitory signaling in NK cells such. Given that exhaustion markers LAG3 and PDCD1 were found to be significantly associated with NK cell cytotoxicity based on the cross-lagged LME model *M*_*NK*_. We model the exhaustion process as:

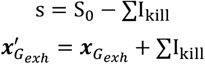

where 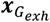 is the vector of RNA expression of exhaustion marker genes *G* = {*LAG*3, *PDCD*1}, which further updates NK cells’ killing probability *p*_*kill*_.

Exhaustion Model 3 (EM3):

Increased expression of exhaustion markers could in turn regulate the synthesis and secretion of cytolytic granules. We used an exponential function to link Hill equations of exhaustion markers to the reduction of serial killing capacity.

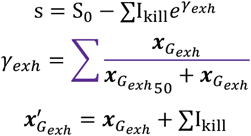

We compared the three EMs by fitting to tumor rechallenge assay in ^5^ and selected the model with the smallest MSE loss. Results are provided in **Supplementary Material 1**.

#### 4.2.3 Vigilant NK cells (*N*_*V*_)

Several studies have demonstrated that NK cells exhibit functions previously only available among cells of the adaptive immune system, including the cell memory^49,70^. Homeostatic proliferation drives NK cells to transform into a dormant, sustained phenotype with preserved effector function and self-renewal potential that can live in the host for prolonged time^48,49,74–79^. To differentiate such NK cells before and after the second viral or pathogen challenge, we termed the former as vigilant NK cells (N_V_) and the latter as memory-like NK cells. In ABMACT, upon local tumor clearance (*B*_−_), cytotoxic NK cells exposed to tumors can transform to vigilant NK cells (N_V_) at a probability *k*_B−_. The vigilant cluster had a lower proliferation rate *p*_*V*_ and death rate *d*_*V*_ to sustain a small population in the host. A proportion of the vigilant NK cells are capable of exhibiting recall upon a second tumor stimulation (event denoted as *B*_+_ when tumor cells are present in the neighborhood) and convert back to cytotoxic NK cells at a probability *k*_*C*+_.

#### 4.2.4 Tumor cells (*B*)

Malignant cells are characterized by uncontrolled growth and immune evasion^80^. B-cell non-Hodgkin lymphoma was characterized to have a medium proliferation rate ranging from 0.15/day to 0.80/day in indolent to highly aggressive types^81^. Glioblastoma was estimated to have a medium proliferation rate of 0.022/day using patient MRI data^82^. The tumor cell agents (e.g. B cell lymphoma, glioblastoma) are modeled to be highly proliferative with a minimum likelihood of apoptosis. As the proliferation rates *p*_*C*_ can be specific to cell lines, host conditions, and other factors, the proliferation rates used in modeling were estimated using tumor-only bio-illuminance data from the lymphoma^23^ and GBM^20^ mouse models (**Supplementary Material 1**). To account for immune evasion, we assumed tumor cells can evade CAR-NKs by downregulating CAR target such as CD19 and gain mutations over generations. We assumed tumor cells has a probability of gaining immune resistant mutation *p*_*mutate*_ of 0.001^83^, which is added to the probability of evading NK cell cytotoxic killing *p*_*evade*_. In models with specific CAR targets such as anti-CD19, the expression of the target in tumor cell agents modifies *p*_*evade*_. For example, *p*_*evade*_ of tumor cells expressing high CD19 proportionally reduce by a constant scaling factor. In addition to proliferation, death, mutation, a tumor cell agent is able to move to neighboring grids at every model step with a probability m_B_.

### 4.3 Integrate functional genetic effects in cell agents

#### 4.3.1 Single-cell RNA sequencing data processing

Longitudinal scRNA-seq data of NK cells and tumor cells were obtained from GSE190976 and processed as described in Supplementary Materials of Li *et al*.^23^. GOBP gene set density scores (GSDS) were calculated using R package “gsdensity”^84^. Pre-infusion scRNA-seq data of IL-15 and IL-21 NK cells^20^ were obtained from GSE227098 in Shanley et al. ^20^.

#### 4.3.2 Feature selection using linear mixed effects modeling

NK cell lysis killing is tightly regulated by a repertoire of activating surface receptors inhibitory receptors such as Fc receptor FcγRIIIa (CD16), NKG2D, and, KIR2DL1^66,69,85^. To understand the molecular features underlying NK cells anti-tumoral capability, we performed feature selection on a literature-curated list of 112 NK cell genes and 5 GO Biological Process (GOBP) pathways^23,69,86–90^ (**Supplementary Materials 2**) that regulate NK cell activation, inhibition, OXPHOS, proliferation, survival, cytotoxicity, regulatory function, and memory function.

To select significant genes and pathways and quantify their effects on NK cell cytotoxicity, we applied linear mixed-effects (LME) modeling to paired scRNA-seq data of NK cells and tumor radiance data from the lymphoma mouse model in Li et al.^23^ using R package lme4^91^. We assume that gene-expression patterns in NK cells are associated with function of NK cells and their anti-tumor control effecting tumor size at the next timepoint, as NK cells required tumor antigen stimulation to sustain and might have drastically waned at the time of sample collection if tumors were cleared. Data at “D28” were excluded due to cell count scarcity. The cross-lagged LME model for NK cells, *M*_*NK*_, included random intercept effects for time and group to consider temporal and inter-group variations. M_n4_ was defined as:

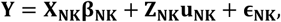

where **Y** = (y_k_)_K_ are means of tumor radiance in unit of p/s/cm^2^/sr across all mice in the k-th group at *t*_2_,…,*t*_*T*_, *k* ∈ {1, …, *K*}. **X** = (x_kig_)_K*I*G_ are gene expressions of the g-th gene of the i-th cell in the k-th group at a timepoint *t*_2_, …, *t*_*T*−1_, *g* ∈ {1, …, *G*_*i*_}, *i* ∈ {1, …, *I*_*k*_}E **β** = (β_g_)_G_ are regression coefficients of fixed effects for the g-th gene. **Z** = (**Z**_**j**_)_J_ are random effects for time and group, *j* ∈ {**t, k**},. **u** = (u_j_)_J_ are random effects regression coefficient for the j-th random effect, 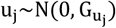 and 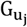 is the covariance matrix, and **ϵ** = (ϵ_ki_)_K*I_: random error for the i-th cell in the k-th group, 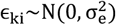. Notations in the format (*X*_*i*_)_*I*_ denotes data *X* of the i-th element in a matrix *I*, and *I* can be multi-dimensional.

As the number of genes and pathways of interest exceeded the number of observations, single covariate LME models were first fitted, and significant covariates were included in the final model. LME models were fitted using restricted maximum likelihood and Nelder-Mead optimizer given small sample sizes. Due to overlaps between genes selected based on literature reviews and relevant GOBP gene sets, we compared models with and without GOBP gene sets and the null model with only random effects using Akaike information criterion (AIC) and Bayesian information criterion (BIC). The LME model with coefficients in **Supplementary Table S2** had the smallest AIC and BIC.

#### 4.3.3 Integrate functional genetic effects in cell agent cytotoxicity

Fixed effects regression coefficients of significant genes and GOBP pathways, **β**_**NK**_, were multiplied with -1 to further parametrize cytotoxic NK cell agents’ probability of killing a tumor cell upon contqact, *p*_*kill*_, as described in Methods section “Cytotoxic killing activity.” In the lymphoma model, the scRNA-seq data at pre-infusion were sampled and randomly assigned to cytotoxic NK cell agents for respective experiment groups. In the GBM model, the scRNA-seq data at pre-infusion were sampled and randomly assigned to cell agents for respective NK cell groups. The expressions multiplied with coefficients derived above and contributed to*p*_*kill*_.

### 4.3 Simulation

#### 4.3.5 Initiation

The simulations were initiated with a 2D discrete grid with 1000 tumor cells and varying NK cells depending on effector-to-target ratio (ETR). Each simulation in lymphoma autonomous growth, rechallenge assay, and mice model simulations was repeated 30 times. Each simulation in the GBM mouse model was repeated 5 times due to computation time constrain. Because of the intra-tumoral administration of NK cells in the GBM mouse model, we assumed no loss of NK cells in infusion and an ETR of 1:5 for all three groups, same as the mouse experiment setup. Simulation data were aggregated to compute the mean and standard deviation.

#### 4.3.5 Model calibration

In lymphoma mouse model, data points at day 0, 7, and 14 post-engraftment (equivalent to day 7 and 14 post-infusion) were used for model calibration. The assumption that engraftment occurred seven days post infusion was based on the comparable tumor radiance in mouse models at day 7 (Figure 2C in Li et al.^23^). In GBM mouse model, data points at day 14, 22, and 37 after intra-tumoral injection were used for model calibration. To reduce the high computation demand of parameter search of ABM, we iteratively optimized parameters from sparse parameter sets to more granular search. The tumor cell proliferation and death rates were fixed before proceeding to the remaining parameters. Mean squared error (MSE) loss was calculated between simulated tumor progression and observed tumor progression obtained from experiment data. We estimated the ETR by selecting the ETR that minimize the total MSE across groups. The remaining parameters were selected by minimizing MSE under the ETR. We evaluated the stability of fitting using coefficients of variations of tumor progression over time.

To benchmark ABMACT, we compare it with the commonly used ordinary differential equation (ODE) models (**Supplementary Material 5**) on the lymphoma mice model dataset. ODE models were constructed with consideration of NK cell cytotoxicity and transitions between phenotypes. ABMACT and ODE models were calibrated on the same xenograft lymphoma mice model data in Li et al. ^23^. The models were evaluated by MSE, and the benchmarking results are provided in **Figure S4b-c**.

### 4.4 Feature importance sensitivity analysis

Aggregating simulation data from in silico perturbation experiments of ABMACT, we trained a Random Forest Regressor (RFR) using scikit-learn^92^ to evaluate the importance of model parameters on accumulated tumor growth and prediction accuracy. Tumor growth was measured by the area under the curve (AUC) over a 35-day simulation period. Prediction accuracy was measured by MSE between simulated data and observed experiment data in the xenograft lymphoma mice model in Li et al.^23^. Feature importance was evaluated by the mean decrease in impurity (MDI) and permutation importance (PI). MDI measures the information gain of features in predicting outcomes. In the case of predicting a continuous outcome variable, MDI measures the reduction in MSE when splitting a variable at a tree node.^93^. PI measures the reduction in the model accuracy score when randomly shuffling a feature’s value, overcoming the potential biases of MDI for highly variable features^94^.

### 4.5 Statistical testing

Two-tailed Mann-Whitney U tests with a significance level of 0.05 were used for testing differences in simulated overall tumor progression (AUC) between conditions. P-value adjustment for multiple testing was performed using Benjamini-Hochberg correction.

### 4.6 Model parameters

The calibrated lymphoma and GBM ABMACT models are initialized with the parameters below in Table 1. A major challenge in ABM is parameter search. We parametrized model and agent properties based on literature where possible, performed calibrations on collected datasets, and conducted local sensitivity analysis on hyperparameters to reduce search space.

**Table 1.** Model parameters.

| Parameter | Description | Value | Unit | Source |
| --- | --- | --- | --- | --- |
| $p_c$ | Proliferation rate of cytotoxic NK cell | Vary by cell product | $day^{-1}$ | Model fitting on data from Liu et al.<sup>19</sup> |
| $d_c$ | Death rate of cytotoxic NK cell | Vary by cell product | $day^{-1}$ | Model fitting on data from Liu et al.<sup>19</sup> |
| $\mu_c$ | Baseline NK cell cytotoxicity mean | Vary by cell product | Dimensionless | Model fitting on data from Li et al.<sup>23</sup> |
| $\sigma_c$ | Baseline NK cell cytotoxicity standard deviation | 0.01 | Dimensionless | Fixed |
| $b$ | NK cell cytotoxicity genetic effect coefficient | 0.1 | Dimensionless | Parameter search |
| $\gamma$ | Chimeric antigen receptor effect exponent | CAR: 0.5<br>NT: 1.0 | Dimensionless | Parameter search |
| $\beta_{G_{NK}}$ | Coefficient vector of significant NK cell cytotoxicity molecular features $G$ | Supplementary Table S2 | Dimensionless | LME model fitting on lymphoma mice model data in Li et al.<sup>23</sup> |
| $g_{NK}$ | Normalized RNA expressions and GOBP densities of NK cells | Supplementary Data 1 | Dimensionless | <sup>23</sup> |
| $m_N$ | Movement probability of NK cells | 0.9 | Dimensionless | Fixed |
| $v_N$ | Movement speed of NK cells | 39 | $\mu m/hr$ | <sup>95</sup> |
| $S_0$ | Initial NK cell serial killing capacity | Vary by cell product | cell | Model fitting on data from Marin et al.<sup>5</sup> |
| $b_{p_c}$ | Natural decay parameter for cytotoxic NK cell proliferation rate | Vary by cell product | $\Delta T^{-1}$ | Model fitting on data from Liu et al.<sup>19</sup> |
| $B_{50}$ | Half-maximum tumor load | 25 | Dimensionless | Model fitting on data from Liu et al.<sup>19</sup> |
| $c_{50}$ | Half-maximum cytokine level | 70 | Dimensionless | Model fitting on data from Liu et al.<sup>19</sup> |
| $\gamma_1$ | Hill's equation exponent of tumor antigen effect | 0.2 | Dimensionless | Model fitting on data from Liu et al.<sup>19</sup> |
| $\gamma_2$ | Hill's equation exponent of cytokine effect | 0.2 | Dimensionless | Model fitting on data from Liu et al.<sup>19</sup> |
| $t_{50_{IL15}}$ | Half-life of IL-15 | 2.5hr | $\Delta T^{-1}$ | <sup>72</sup> |
| $t_{50_{IL21}}$ | Half-life of IL-21 | 0.2hr | $\Delta T^{-1}$ | <sup>96</sup> |
| $p_{evade}$ | Probability of B cell lymphoma tumor cells evading immune surveillance | $\propto g_{\{B,CD19\}}^{-1}$ | Dimensionless | <sup>97</sup> |
| $p_V$ | Proliferation rate of vigilant NK cells | 1e-3 | $day^{-1}$ | Fixed to be a small value based on literature 48,49,74–79 |
| $d_V$ | Death rate of vigilant NK cells | 1e-3 | $day^{-1}$ | Fixed to be a small value based on literature 48,49,74–79 |
| $p_E$ | Proliferation rate of exhausted NK cells | 0 | $day^{-1}$ | Fixed |
| $d_E$ | Death rate of exhausted NK cells | Vary by cell product | $day^{-1}$ | Model fitting on data from Liu et al.<sup>19</sup> |
| $k_{B-}$ | Transition probability of $N_C$ to $N_V$ | 1.0 | Dimensionless | Fixed |
| $k_{B+}$ | Transition probability of $N_V$ to $N_C$ | 0.9 | Dimensionless | Fixed |
| $p_{B_{lymphoma}}$ | Proliferation rate of B cell lymphoma tumor cell | 0.455 | $day^{-1}$ | Modeling fitting on data from Li et al.<sup>23</sup> |
| $p_{B_{glioblastoma}}$ | Proliferation rate of glioblastoma tumor cell | 0.223 | $day^{-1}$ | Modeling fitting on data from Shanley et al.<sup>20</sup> |
| $d_B$ | Death rate of tumor cell | 1e-4 | $day^{-1}$ | Modeling fitting on data from Li et al.<sup>23</sup> |
| $m_B$ | Movement probability of tumor cells | 0.1 | Dimensionless | Fixed |
| $v_B$ | Movement speed of tumor cells | 1 | $grid/step$ | Fixed |
| $p_{mutate}$ | Percentage increase of $p_{evade}$ | 1e-3 | Dimensionless | Fixed |
| $\Delta T$ | Model step length | 4 | $hr$ | <sup>66</sup> |
| $l$ | Grid cell size | 50 | $\mu m$ | Fixed |
| $B_{Max}$ | Maximum number of tumor cells per grid cell | 25 | cell | Fixed |

#### Software

The feature selection using LME model was performed in R (v4.2.2) ^98^. ABM simulations were performed in Python (v3.9) ^99^ using the MESA framework^63^.

## Code Availability

The source code for reproducing the work is accessible at: https://github.com/KChen-lab/ABMACT, upon acceptance of the publication. Private link for reviewers: https://zenodo.org/records/14884628?token=eyJhbGciOiJIUzUxMiJ9.eyJpZCI6IjI1NmM0YzNjLWM4NDItNGZhMy1iNDIxLWExNDBmZGIxZjJhZiIsImRhdGEiOnt9LCJyYW5kb20iOiIyNGZjNWIxYzAwM2NiZGI2Njk2NWIyMzg4ZGQwNGRmOCJ9.JXMZmpYk113L3×3cgEPwZ-GPKZCsHMA7aHtLeAOHR2hzx5lZngmIJRvKIdgMfGWF4sy01rAFHpuTxTxAHlKm6g

## Acknowledgements

This work is made possible by 2024-345892 from the Chan Zuckerberg Initiative DAF, an advised fund of the Chan Zuckerberg Initiative Foundation, 5U01CA281902 from National Cancer Institute, and 75N99223S0001 from the Advanced Research Projects Agency for Health (ARPA-H). This work was supported in part by the University of Texas MD Anderson Cancer Center Institute for Cell Therapy Discovery & Innovation. The data used in this study were supported, in part, by grants (1 R01 CA211044-01, 5 P01CA148600-03, and U01CA247760) from the National Institutes of Health (NIH), the Cancer Prevention and Research Institute of Texas (grants RP180466 and RP180248), the Leukemia Specialized Program of Research Excellence (SPORE) Grant (P50CA100632), the Specialized Program of Research Excellence (SPORE) in Brain Cancer grant (P50CA127001), and CPRIT Single Core (RP180684), and a grant (P30 CA016672) from the NIH to the MD Anderson Cancer Center Flow Cytometry and Cellular Imaging Core Facility that assisted with the mass cytometry studies. We sincerely thank Drs. Eleonora Dondossola, Peng Wei, Ziyi Li, Enli Liu, and Li Li for their support and insightful feedback during the development of the study.

