## Supplementary Materials for "Agent-based modeling of cellular dynamics in adoptive cell therapy"

Supplementary Figures

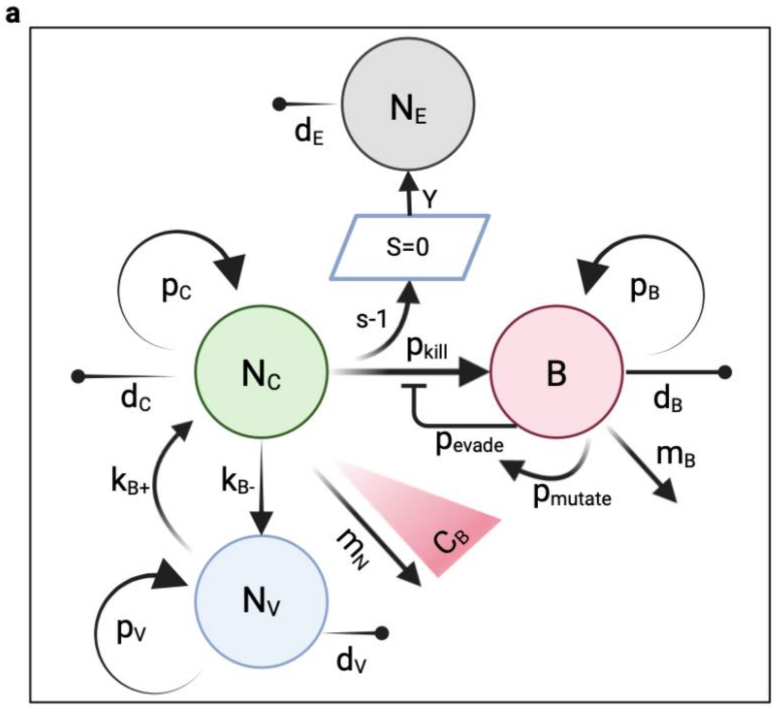


Figure S1: ABMACT model structure, comprising three populations of NK cells – cytotoxic NK cells ($N_c$), exhausted NK cells ($N_E$), vigilant NK cells ($N_V$) – and tumor cells ($B$). The model was built using MESA [1], an open-source Python library. $N_C$: cytotoxic NK cells. $N_V$: vigilant NK cells. $N_E$: exhausted NK cells. $B$: B cell lyphoma tumor cells. $p_{kill}$: probability of NK cells killing tumors upon contact. $p_{evade}$: probability of tumor cells evading immune surveillance. $p_{mutate}$: probability of tumor cells gaining anti-immune mutations. $s$: NK cell serial killing capacity, which reduces the number of killing till exhaustion ($s = 0$). $p_C$, $p_V$, $p_B$: proliferation probability of $N_C$, $N_V$, and $B$, respectively. $d_C$, $d_E$, $d_V$, $d_B$: apoptosis probability of $N_C$, $N_E$, $N_V$, and $B$, respectively. $k_{B-}$, $k_{B+}$: probability of $N_c$ transform to $N_V$ upon tumor clearance and the reverse upon a second stimulation. $m_N$, $m_B$: probability of locomotion of NK and tumor cells; NK cells move against tumor antigen gradient $c_B$.

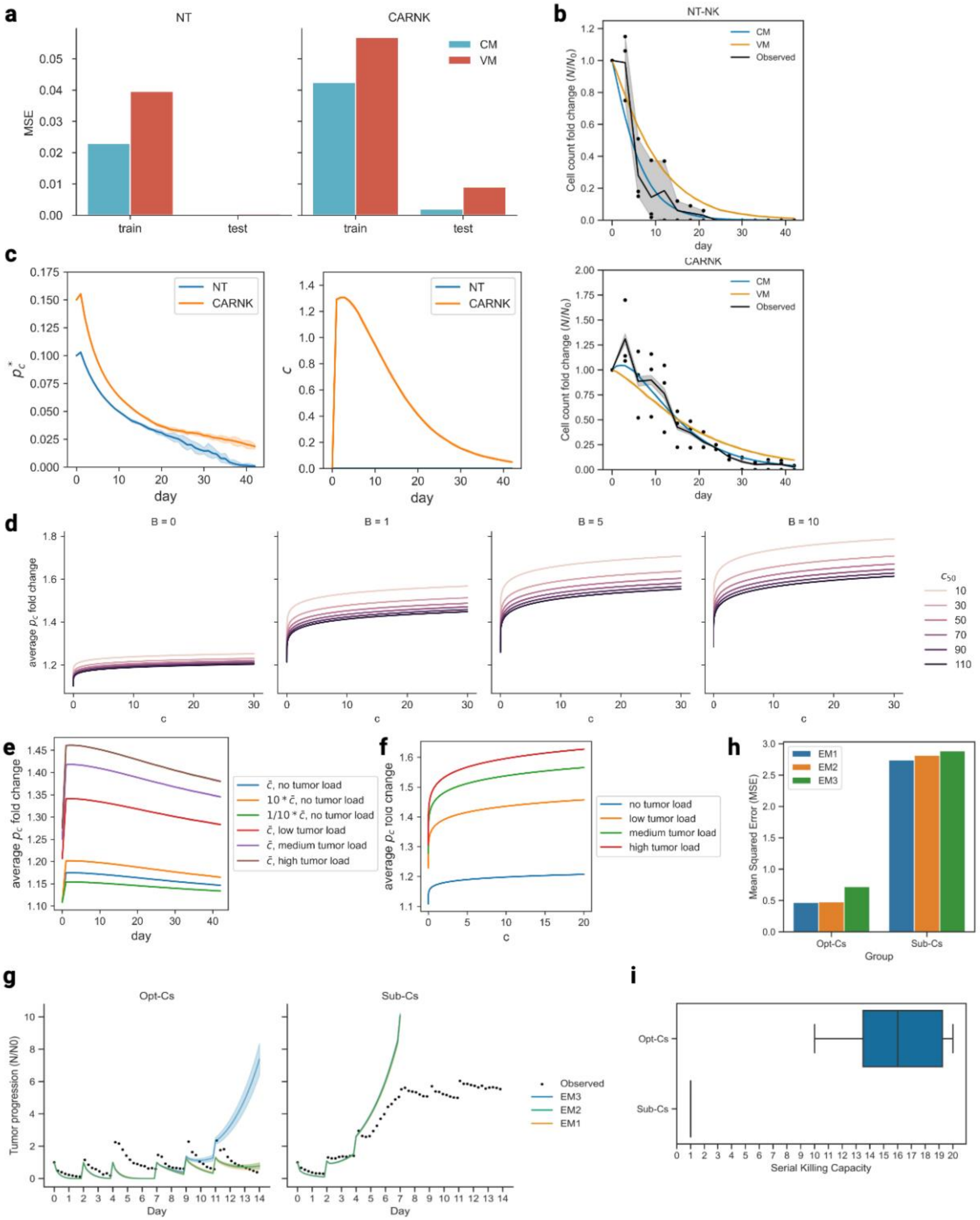


Figure S2: Initial estimation of NK cell properties and characterization of NK cell functions: NK cell proliferation: (a) Fitted population fold changes of NT-NKs and CD19IL15 CAR-NKs in autonomous growth assay[2] using ABM with two NK cell proliferation models. CM: cytokine-dependent model. VM: vanilla model. Cell count fold change calculated by normalizing cell count ($N$) with respect to initial cell count ($N_0$). (b) Train and test mean squared error (MSE) of NT-NKs and CD19IL15 CAR-NKs. The first

70% of data were used for fitting and the remaining for testing. (c) Average proliferation rate (left) and cytokine level (right) of NT-NKs and CD19IL15 CAR-NKs simulated by ABM with CM. (d) Simulated average proliferation rate $p_c$ fold change with respect to cytokine level at different tumor load ($B$, increasing horizontally) and sensitivity to cytokine stimulation ($c_{50}$, half-maximum cytokine level, smaller value indicates higher sensitivity). Fold change calculated by average $p_c$ with respect to $p_c$ at $c = 0$. (e) Simulated average proliferation rate fold change over time at different tumor load and sensitivity to cytokine stimulation. $\bar{c}$: mean half-maximum cytokine level. Fold change calculated by average $p_c$ with respect to $p_c$ at $t = 0$. NK cell cytotoxicity (f) Fitted dose-response curves for CD19IL15 CAR-NKs, CD19 CAR-NKs, and NT-NKs cocultured with Raji lymphoma cell line [3] using ordinary differential equations (ODE). Cytotoxicity level measured by Cr-release assay. NK cell exhaustion: (j) Fitted tumor population fold change of optimal cord CD19IL15 CAR-NKs (Opt-Cs) and suboptimal cord CD19IL15 CAR-NKs (Sub-Cs) in Raji lymphoma rechallenge assay[4] using ABM with three exhaustion models (EM). (h) MSE of three Ems for Opt-Cs and Sub-Cs. All data are used for fitting as splitting to train and test led to large deviation in late phase of rechallenge. (i) Estimated serial killing capacity of Opt-Cs and Sub-Cs in fitted simulations.

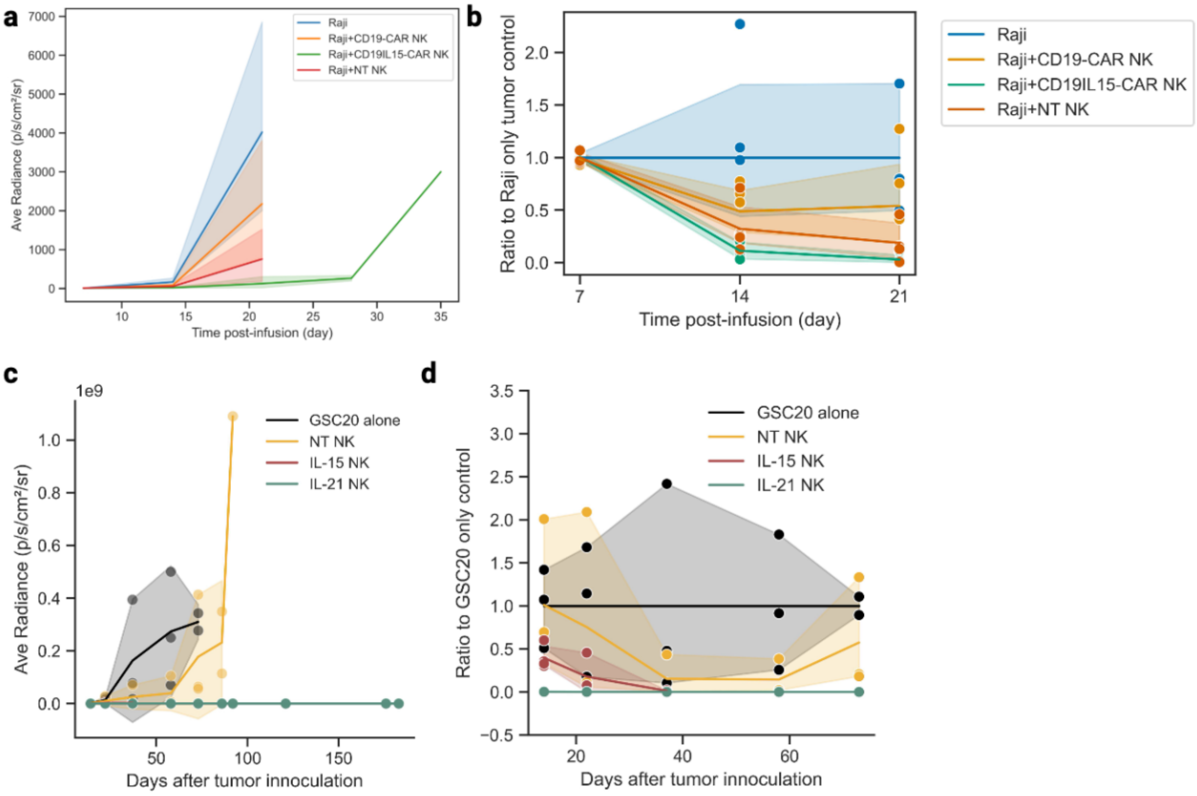


Figure S3: Lymphoma[5] and glioblastoma[6] mouse models. (a) Tumor radiance over time post-infusion of NK cells. (b) Data used for characterizing ABMACT lymphoma mouse model. Normalized tumor progression over time post-engraftment of tumor and NK cells. Engraftment was assumed to be at day 7 post-infusion, given comparable tumor radiance at day 7 post-infusion. Normalized tumor progression is calculated by dividing tumor radiance in treatment groups by radiance in control group at each timepoint. (c) Tumor radiance over time post tumor inoculation. (d) Data used for characterizing ABMACT glioblastoma mouse model. Normalized tumor progression over time post tumor inoculation up to day 37.

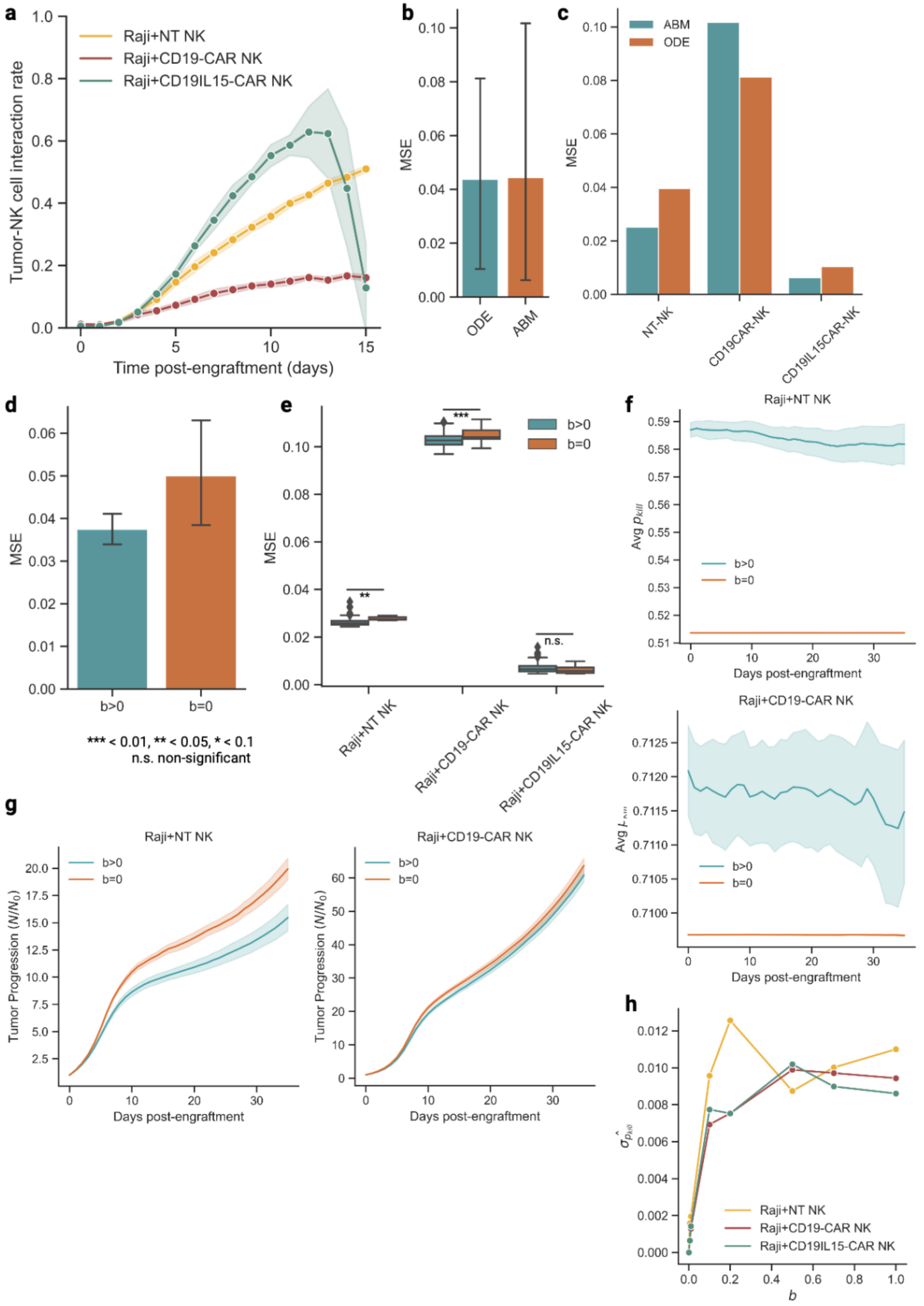
a
Raji+NT NK
Raji+CD19-CAR NK
Raji+CD19IL15-CAR NK
Tumor-NK cell interaction rate
Time post-engraftment (days)
b
MSE
ODE
ABM
c
ABM
ODE
MSE
NT-NK
CD19CAR-NK
CD19IL15CAR-NK
d
MSE
b>0
b=0
*** < 0.01, ** < 0.05, * < 0.1
n.s. non-significant
e
b>0
b=0
MSE
***
**
n.s.
Raji+NT NK
Raji+CD19-CAR NK
Raji+CD19IL15-CAR NK
f
Raji+NT NK
b>0
b=0
Days post-engraftment
Raji+CD19-CAR NK
b>0
b=0
Days post-engraftment
g
Raji+NT NK
b>0
b=0
Tumor Progression (N/N0)
Days post-engraftment
Raji+CD19-CAR NK
b>0
b=0
Tumor Progression (N/N0)
Days post-engraftment
h
Raji+NT NK
Raji+CD19-CAR NK
Raji+CD19IL15-CAR NK
b

Figure S4: Benchmark ABMACT model fitting and genetic effects. (a) Interaction rate between tumor cells and cytotoxic NK cells in the lymphoma mouse model. Interaction rate calculated by the ratio between tumor cells co-locating with cytotoxic NK cells and the total tumor cell count. (b) Overall MSE comparison between ABMACT and ordinary differential equations (ODE) method in lymphoma mouse model simulation and (c) by group. (d) Overall MSE comparison between the lymphoma model including genetic effects ($b > 0$) and without ($b = 0$) and (e) by group. NK cell cytotoxicity genetic effect coefficient $b$ was varied to compare MSE. $b \in \{0.0, 0.005, 0.01, 0.10, 0.2, 0.5, 0.7, 1.0\}$. (f) Distribution of NK cytotoxic killing probability $p_{kill}$ with ($b > 0$) and without ($b = 0$) genetic effects. (g) Tumor progression over time with ($b > 0$) and without ($b = 0$) genetic effects in two groups with significant MSE improvement by including genetic effects. Left: NT-NK cell group. Right: CD19 CAR-NK cell group. (h) Standard deviations of NK cytotoxic killing probability, $\hat{\sigma}_{p_{kill}}$, increase with $b$. Interval bands of 2 s.e. were calculated using a bootstrap of 1000 iterations.

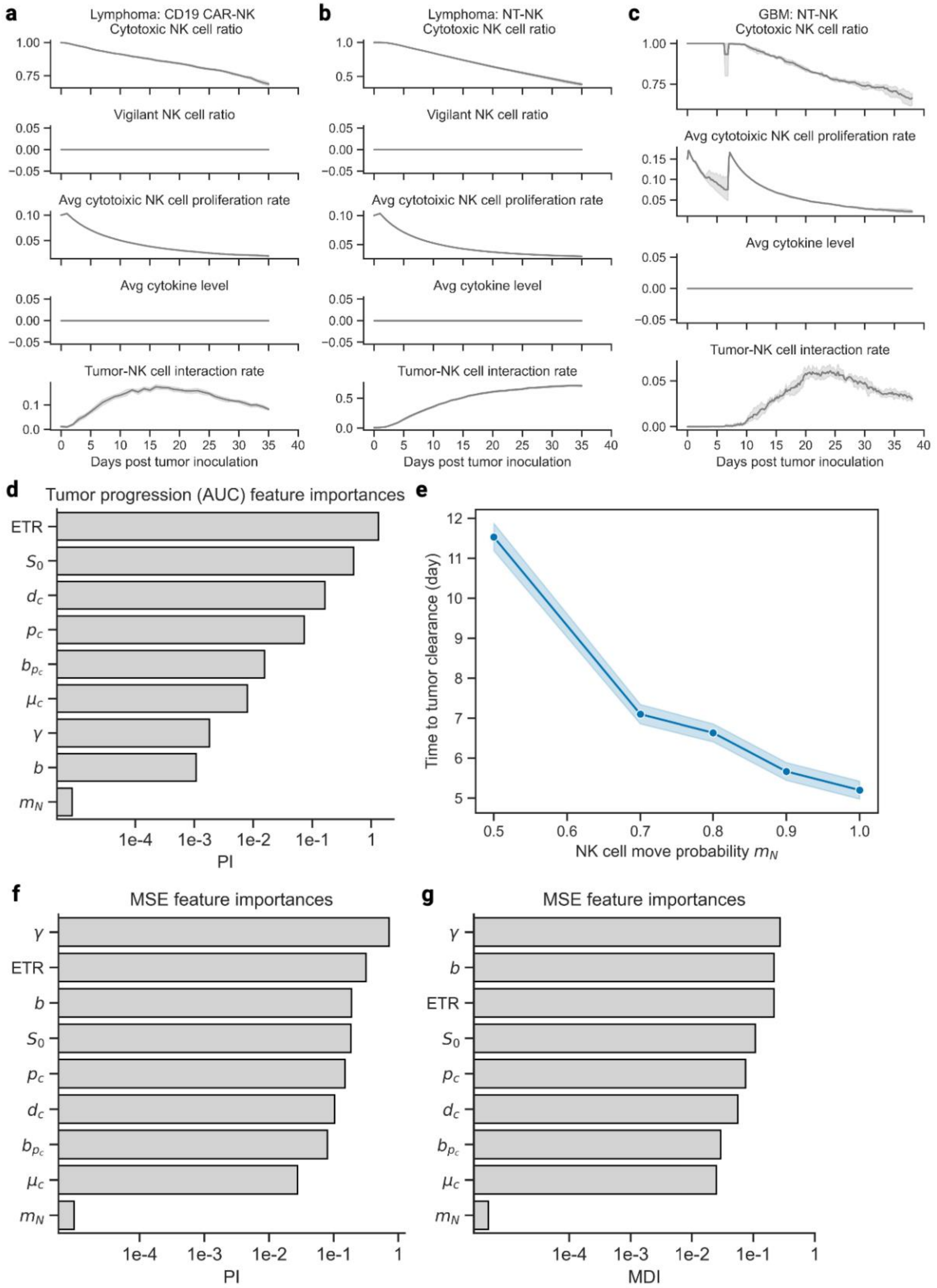
a
Lymphoma: CD19 CAR-NK
Cytotoxic NK cell ratio
Vigilant NK cell ratio
Avg cytotoixic NK cell proliferation rate
Avg cytokine level
Tumor-NK cell interaction rate
Days post tumor inoculation
b
Lymphoma: NT-NK
Cytotoxic NK cell ratio
Vigilant NK cell ratio
Avg cytotoixic NK cell proliferation rate
Avg cytokine level
Tumor-NK cell interaction rate
Days post tumor inoculation
c
GBM: NT-NK
Cytotoxic NK cell ratio
Avg cytotoixic NK cell proliferation rate
Avg cytokine level
Tumor-NK cell interaction rate
Days post tumor inoculation
d
Tumor progression (AUC) feature importances
ETR
$S_0$
$d_c$
$p_c$
$b_{p_c}$
$\mu_c$
$\gamma$
$b$
$m_N$
1e-4 1e-3 1e-2 1e-1 1
PI
e
Time to tumor clearance (day)
NK cell move probability $m_N$
f
MSE feature importances
$\gamma$
ETR
$b$
$S_0$
$p_c$
$d_c$
$b_{p_c}$
$\mu_c$
$m_N$
1e-4 1e-3 1e-2 1e-1 1
PI
g
MSE feature importances
$\gamma$
$b$
ETR
$S_0$
$p_c$
$d_c$
$b_{p_c}$
$\mu_c$
$m_N$
1e-4 1e-3 1e-2 1e-1 1
MDI

Figure S5: Cytotoxic NK cell ratio, vigilant NK cell ratio, average cytotoxic NK cell proliferation rate, average cytokine level, and tumor-NK cell interaction rate of (a) CD19 CAR-NK cells in the lymphoma model, (b) NT-NK cells in the lymphoma model, and (c) NT-NK cells in the GBM model. NK cell subtype ratio calculated with respect to the total NK cell population at each timepoint. Tumor-NK interaction rate calculated by the ratio of tumor cells with co-locating cytotoxic NK cells with respect to the total tumor cell population. Sensitivity analysis of ABMACT model parameters. (d) Random Forest regression feature importance of model parameters on tumor progression area under the curve (AUC) measured by permutation index (PI) (**Methods**). (e) Time to tumor clearance by cytotoxic NK cell movement probability $m_N$. Random Forest regression feature importance of model parameters on MSE measured by (f) mean decrease in impurity (MDI) and (g) PI.

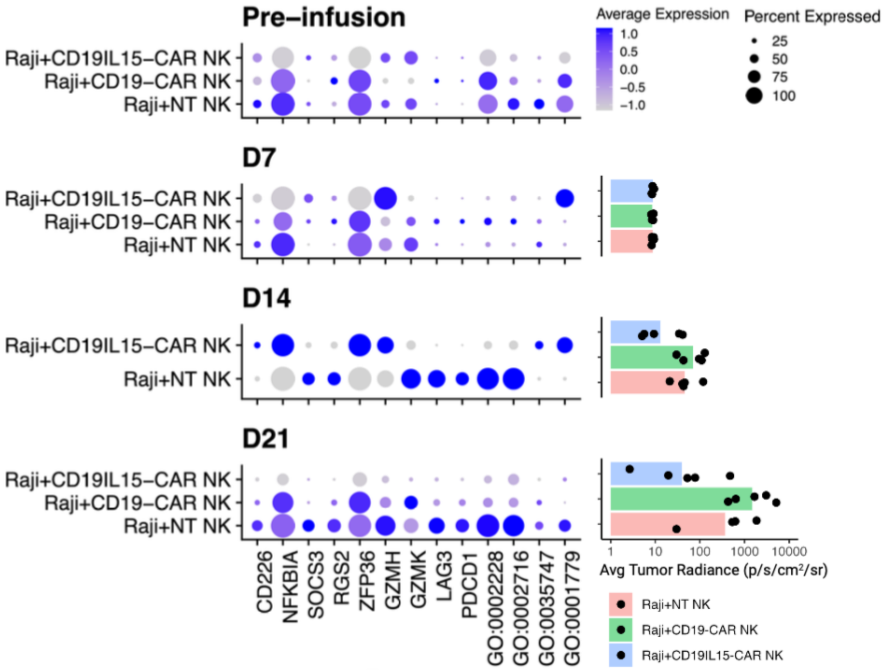


Figure S6: Expressions of genes and pathways selected by LME model and tumor loads over time. Tumor loads were measured by bio-illuminance radiance in Raji lymphoma PDX mice model (n=5 per group)[5]. Average expressions represented by the color scale with circle sizes corresponding to percentages of cells expressing the genes or pathways. Average tumor radiance by group (bar) and replicate (dot) on the right.

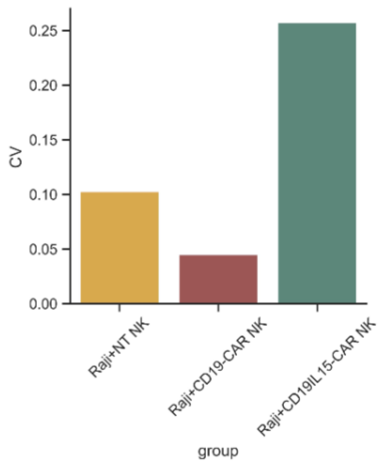


Figure S7: Coefficient of variation ($CV = \sigma/\mu$) on fitted simulation results of the lymphoma model.

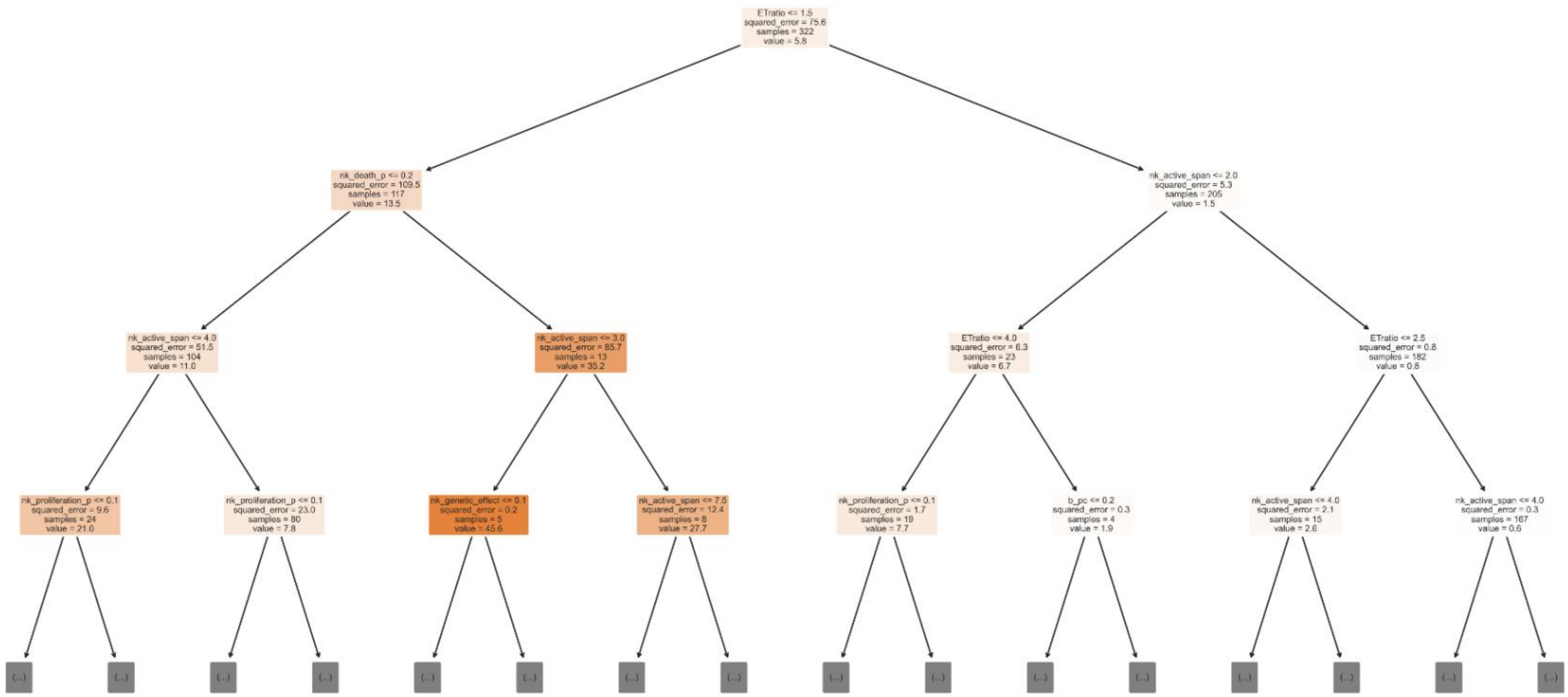

ETratio <= 1.5
squared_error = 75.6
samples = 322
value = 5.8
nk_death_p <= 0.2
squared_error = 109.5
samples = 117
value = 13.5
nk_active_span <= 2.0
squared_error = 5.3
samples = 205
value = 1.5
nk_active_span <= 4.0
squared_error = 51.5
samples = 104
value = 11.0
nk_active_span <= 3.0
squared_error = 85.7
samples = 13
value = 35.2
ETratio <= 4.0
squared_error = 6.3
samples = 23
value = 6.7
ETratio <= 2.5
squared_error = 0.8
samples = 182
value = 0.8
nk_proliferation_p <= 0.1
squared_error = 9.6
samples = 24
value = 21.0
nk_proliferation_p <= 0.1
squared_error = 23.0
samples = 80
value = 7.8
nk_genetic_effect <= 0.1
squared_error = 0.2
samples = 5
value = 45.6
nk_active_span <= 7.5
squared_error = 12.4
samples = 8
value = 27.7
nk_proliferation_p <= 0.1
squared_error = 1.7
samples = 19
value = 7.7
b_pc <= 0.2
squared_error = 0.3
samples = 4
value = 1.9
nk_active_span <= 4.0
squared_error = 2.1
samples = 15
value = 2.6
nk_active_span <= 4.0
squared_error = 0.3
samples = 167
value = 0.6

Figure S8: Top three splitting of model parameters of the Random Forests Regressor in sensitivity analysis.

Supplementary Materials

### 1 Cell agents design

Given the large search space of numerous parameters in agent-based models (ABM), we perform initial parameter estimations and characterize mathematical representations of cell functions based on existing experiment data. While cellular processes such as proliferation, exhaustion, death, antigen recognition, and migration are well established by probability distributions in prior modeling works[7–10], some of key processes governing cytotoxic NK cell kinetics and life cycles can be more complex to model. In view of data availability, modeling accuracy, and computational tractability, we further parameterized these agents using data obtained from in vitro autonomous growth and rechallenge assays (**Methods**). In the following sections, we outline pre-ABM estimations for NK cell proliferation, cytotoxicity, exhaustion, and tumor cell proliferation.

#### 1.1 Characterize NK cell proliferation kinetics

NK cell proliferation and persistence after infusion to patients are critical to treatment response. ACT NK cell products are often transfected with cytokine-expressing vectors to boost their expansion after infusion. To characterize NK cell proliferative capability, we constructed mathematical models and fitted to cell-line autonomous growth experiment data. We hypothesized that NK cell proliferation dynamically changes with cytokine stimulation and compared a cytokine-dependent model (CM) and a vanilla model (VM) with a fixed decay rate and no cytokine dependence (**Methods**). We integrated CM and VM in proliferation function of cytotoxic NK cell agents in ABMACT and evaluated fitting to NK cell autonomous growth assay[2] using mean squared error (MSE). The fitting results are provided in **Table S1**. Splitting the first 70% of data for training and the remaining for testing, ABM with CM resulted in lower MSE than VM in both CAR-NKs and NT-NKs (**Figure S2a**). Cell count fold changes in both NT-NKs and CAR-NKs were captured more closely using CM than VM (**Figure S2b**). CAR-NKs exhibited higher proliferation rate over time and expressed cytokines (**Figure S2d**).

In addition to cytokine level, CM considers the effect of tumor antigen stimulation, represented by tumor load $B$, and sensitivity to cytokine, represented by half-maximum level of cytokine $c_{50}$. Increasing cytokine level, sensitivity to cytokine (smaller $c_{50}$), and tumor load resulted in higher proliferation (**Figure S2d-f**). Under varying levels of tumor load $B$, the proliferation modulating effect displayed similar trends across sensitivity to cytokine stimulation (half-maximum cytokine level $c_{50}$). Without tumor load, the proliferate rate fold change was in the range of [1.10, 1.25], while with tumor load, it was in the range of [1.21, 1.79]. Notably, even with low tumor load, the modulating effect is stronger than increasing cytokine level or sensitivity at B=0 (p-val<1e-4, Wilcox Rank Sum test). This corresponded with prior findings that NK cells with IL-15 alone do not result in uncontrolled growth and require presence of pathogens or antigens to expand[2]. NK cells can be engineered to express IL-15R$\alpha$ to increase their sensitivity to endogenous IL-15[11]. We parametrized the sensitivity to IL-15 as the half-maximum cytokine level $c_{50}$ and compared the modulating effect by varying $c_{50}$ (**Figure S2d**). Even by reducing $c_{50}$ from the fitted value of 70 to 50, the modulating effect on NK cell proliferation significantly improved (p-val<1e-4, Wilcox Rank Sum test). The simulated results suggest that high concentrations of cytokines may not be required to significantly boost NK cells' proliferation capacity. Engineered NK cells can achieve optimal benefits at moderate cytokine levels by adequate transduction rates of cytokine-expressing vectors. While high cytokine concentrations may elevate the risk of cytokine-induced toxicity or activation-induced cell death, they do not proportionally enhance NK cell expansion capabilities. As hyperparameters for the tumor antigen effect could not be validated using the same dataset, we performed sensitivity analysis by varying the half-maximum tumor load $B_{50}$ and Hill equation exponent $\gamma_1$ and confirmed the same observation.

Table S1 Fitted parameters of cytokine-dependent (CM) and vanilla model (VM) for NK cell proliferation.

| | *Cytokine-dependent model* | | *Vanilla model* | |
|---|---|---|---|---|
| | CARNK | NT | CARNK | NT |
| $p_c$ | 0.15 | 0.1 | 0.05 | 0.05 |
| $d$ | 0.10 | 0.20 | 0.05 | 0.10 |
| $c_{50}$ | 70 | / | / | / |
| $\gamma_2$ | 0.2 | / | / | / |
| $b$ | 0.19 | 0.13 | 0.10 | 0.10 |

#### 1.2 Initial estimation of NK cell baseline cytotoxicity

To reduce parameter search space for NK cell cytotoxicity in ABM, we performed initial estimation of relative baseline cytotoxicity of CAR19IL15 NK cells, CAR19 NK cells, and NT NK cells using the dose-response data from the $^{51}$Cr-release assay of NK cell products against Raji targets in Li et al.[3]. We applied the following dose-response Emax model:

$$E(y) = \frac{E_0}{1 + \left(\frac{R * x_g}{k_{50}}\right)^m}$$

Where $x_g$ denoted the average relative NK cell cytotoxicity of the group g, $E_0$ denoted the baseline death percentage (cytotoxicity) of tumor cells at ratio = 0, R denoted the effector:target ratio, $k_{50_g}$ denoted the half-maximum relative average NK cell cytotoxicity of the group g, and m denoted Hill's coefficient (shape parameter). We assumed $E_0$ is constant across groups given the same Raji tumor cells used in experiments. The nonlinear least squares estimation was performed using the least_squares function of the Python package scipy[12]. Three types of loss including "linear", "soft_l1", and "cauchy" were compared to select parameters with minimum loss. The best fit was found using the linear loss. The relative cytotoxicity of CAR19IL15 NK cells, CAR19 NK cells, and NT NK cells were estimated to be 0.271, 0.016, 0.002, respectively.

The relative cytotoxicity of engineered NK cells against GSC20 glioblastoma cell line were estimated from real-time killing assay[6]. The estimations were obtained from mean percentage of GSC20 killing after killing saturated (hr>26). The relative cytotoxicity of IL-21NKs, IL-15 NKs, and NT-NKs were estimated to be 0.515, 0.629, 0.401, respectively.

While the estimations of relative cytotoxicity do not directly translate to probability of killing, they serve as priors in subsequent ABM fitting to reduce search space.

#### 1.3 Characterize NK cell exhaustion process

NK cell exhaustion was found to be associated with decreased degranulation of cytolytic molecules, impaired cytotoxicity towards target cells, dysregulated inhibitory and activating signaling, and suppressed metabolism[13,14]. NK cells' ability to degranulate is determined by the number of cytolytic granules stored in cells[15] as well as tight regulations by a repertoire of activation and inhibitory signals[16,17]. Exhaustion in NK cells impair their immunosurveillance function and leads to ineffective tumor control[13]. Conventionally, NK cell exhaustion is functionally characterized by rechallenge assay. When NK cells lose ability to control tumor growth after repeated tumor cell infusion, the NK cells are deemed to be exhausted. Albeit vigorous research on NK cells, the exact mechanisms underlying the exhaustion process are not fully understood or evaluated quantitatively. This motivated us to evaluate this critical process in NK cell immunity through mathematical modeling embedded in ABMACT.

In this study, we focused on the aspect of exhaustion that directly impacts tumor killing. Gwalani et al.[15] proposed that the efficiency of NK cells in killing target cells was influenced by the number of degranulation, the size and content of lytic granules, and the speed at which these granules are released at the immunological synapse. In the same study, LAMP-1-pHluorin transduced human NK cell lines YTS and NK92 cells were measured to contain $194 \pm 51$ and $206 \pm 57$ perforin positive lytic granules on average using live cell imaging assays, in which single degranulation event released $16 \pm 8$ and $12 \pm 7$ lytic granules, respectively, although merely 2 to 4 degranulation were required to cause effect target cell death[15]. Considering that lytic granules are constitutively expressed in NK cells and pre-formed without requiring stimulations, it is reasonable to assume that each NK cell is capable of initiating multiple times of degranulation in tumor killing. Therefore, we parametrized the NK cell serial killing capacity $s$ to serve as a surrogate for NK cells' degranulation and quality of lytic granules, reflecting their long-term killing effectiveness.

NK cell cytotoxicity towards tumor cells is tightly regulated by a repertoire of activating and inhibitory signals. Several studies reported the association between the expression of PD-1 and LAG-3, which could be induced by interactions with tumor cells, and dysfunction of NK cell cytotoxicity[16,17]. Analyzing longitudinal RNA expressions of CAR-NK cells and tumor radiance in mouse model experiment in Raji lymphoma treated with CAR-NK cells, we found that the expression of NK cell exhaustion markers LAG-3 and PDCD-1 were significantly associated with NK cells' tumor control capability (**Figure 2a**). Therefore, we hypothesized that in addition to reduced degranulation, NK cell exhaustion could also be characterized by reduced killing ability through the increase of inhibitory marker gene expression such as PDCD-1 and LAG-3.

We proposed three mathematical models to describe the NK cell exhaustion process involving reductions in serial killing capacity (SKC) and impaired cytolytic killing induced by exhaustion marker signaling. We evaluated the models using longitudinally measured tumor cell index in rechallenge assays of cord blood (CB) CD19IL15 CAR-NK cells [7] (**Methods**). We found that Exhaustion Model 1 (EM1), where NK cells' SKC linearly reduced with the number of tumor cells killed, recapitulated the experimental observation of CB CD19IL CAR-NK cell rechallenge assay [4] with the highest accuracy among the three models (**Figure S2g-h**). The simulations found that the suboptimal cord cohort (Sub-Cs) exhausted immediately after one kill, while the SKC for Opt-Cs ranged from 10 to 20 (**Figure S2i**), suggesting the contribution of high SKC to superior tumor control ability. Additionally, we observed in the optimal cord cohort (Opt-Cs) that Exhaustion Model 2 (EM2), which modeled the impaired cytotoxicity in addition to linear reduction in SKC, had comparable performance with EM1, suggesting the possibility of tumor killing induced cytotoxicity reduction. Overall, integrating the exhaustion submodule with ABMACT enables the model to capture periodical variations of tumor loads in rechallenge assay and allow us to investigate underlying functional changes leading to the differential tumor control ability.

#### 1.4 Estimate tumor cell proliferation rate

A hallmark of cancer is uncontrolled proliferation[18].Bio-illuminance imaging data of lymphoma[3] and glioblastoma (GBM)[6] mouse models both showed exponential growth in tumor-only group. Given the relatively simpler kinetics as compared to NK cells, we estimated proliferation rate of Raji lymphoma and GSC20 glioblastoma tumor cells using the following ODE model:

$$\frac{dB}{dt} = \mu\left(1 - \frac{B}{B_{max}}\right)B$$

Where B is average tumor radiance, which is a surrogate for tumor size, μ is tumor cell proliferate rate in day$^{-1}$, and $B_{max}$ is the equivalent average radiance of growth-limiting tumor size.

In lymphoma model, the proliferation rate of Raji lymphoma tumor cells was estimated to be 0.444/day using tumor radiance of Raji-only lymphoma mouse model in Li et al.[3]. The best fit was found using lmfit minimize function and Broyden–Fletcher–Goldfarb–Shanno (BFGS) optimizer.

In GBM model, the proliferation rate of GSC20 GBM tumor cells was estimated to be 0.224/day using tumor radiance of GSC20-only GBM mouse model in [6]. The best fit was found using lmfit minimize function and least squares optimizer.

#### 1.5 TME specifications

The simulations are initialized in a 2D Moore space of $50 \times 50$ grids. Each cell agent is assumed to have an average diameter of $10\mu m$ [10]. Each grid is assumed to be $50\mu m \times 50\mu m$ and can hold up to 25 cells. Cytokine molecules diffuse on the matrix using a Gaussian filter and degrade over time. The degradation rate of IL-15 and IL-21 are specified in the parameter table in Methods. Both IL-15 and Il-21 have 162 amino acids[19]. A ribosome in a cell on average can synthesize 162 amino acids in 2 minutes and there are hundreds to millions of ribosomes in a cell[20]. Ribosomal activities vary between cell and cell status. The number of IL-15 or IL-21 that an NK cell secrete per hour can range from 45 to 45 million. For ease of calculation, we assumed that a cytokine-carrying NK cell can secrete 10 units of IL-15 or IL-21 in every model step and calibrated cells' sensitivity to the molecules for model fitting. In every model step, all cell agents with "alive" status are randomly looped through to perform cellular functions and interact.

### 2 Molecular Feature Selection

#### 2.1 NK cell cytotoxicity model $M_{NK}$

We collected a literature-curated list of 112 NK cell genes and 5 GO Biological Process (GOBP) pathways [3,13,21–25] that regulate NK cell activation, inhibition, OXPHOS, proliferation, survival, cytotoxicity, regulatory function, and memory function: CD226, CD244, CD69, FCGR3A, GNLY, IFNG, IL2RA, KIR2DS1, KIR2DS2, KIR2DS4, KIR3DS1, KIT, KLRC2, KLRK1, LAMP1, NCAM1, NCR1, NCR2, NCR3, AKT1, AKT2, AKT3, DUSP1, IER2, JAK1, JAK2, JAK3, MTOR, NFKBIA, NFKBIZ, PTEN, RICTOR, RPTOR, SOCS1, SOCS3, STAT1, STAT2, STAT3, STAT4, STAT5A, STAT6, BAG3, CALU, DNAJB1, HSP90AA1, HSP90AB1, HSP90B1, HSPA1A, HSPA1B, HSPA6, HSPB1, HSPH1, RGS1, RGS2, SLC2A3, UBC, ZFAND2A, ZFP36, ZFP36L1, CCL2, CCL3, CCL4, CCL5, CXCR3, IL15, IL18, IL1B, IL6, IL7, CD247, FASLG, GZMA, GZMB, GZMH, GZMK, GZMM, PRF1, SYK, TNFSF10, ZAP70, HAVCR2, LAG3, TIGIT, HK2, KIR2DL1, KIR2DL3, KIR2DL4, KIR3DL1, KIR3DL2, KLRB1, KLRC1, KLRD1, LILRB1, PDCD1, SIGLEC7, FOXP3, EGR1, EOMES, FOS, FOSB, HIF1A, JUN, JUNB, MYC, NR4A1, NR4A2, NR4A3, TBX21, MAX, MNT, MXD1, MXD4,

GOBP_NATURAL_KILLER_CELL_ACTIVATION,
GOBP_NATURAL_KILLER_CELL_MEDIATED_IMMUNITY,
GOBP_NEGATIVE_REGULATION_OF_NATURAL_KILLER_CELL_MEDIATED_IMMUNITY,
GOBP_NATURAL_KILLER_CELL_CHEMOTAXIS,
GOBP_NATURAL_KILLER_CELL_DIFFERENTIATION.

To select significant genes and pathways and quantify their effects on NK cell cytotoxicity, we applied a cross-lagged linear mixed-effects (LME) model, $M_{NK}$, to paired scRNA-seq data of NK cells and tumor radiance data from the lymphoma mouse model[5] using R package lme4[26]. The derived coefficients of selected genes and pathways were used in cytotoxic NK cell agents' killing probability function.

We proposed that tumor control effect was associated with NK cell scRNA-seq expression in the previous time point, as NK cells required tumor antigen stimulation to sustain and might have drastically waned at the time of sample collection if tumors were cleared. The cross-lagged LME model accounted for the temporal delay between NK cell activity and observable tumor progression or regression, allowing us to identify key genes and pathways contributing to variations in tumor control. For instance, GZMH and GOBP pathway for NK cell differentiation (GO:0001779) showed higher expression in the CD19IL15-CARNK group compared to CD19 and NT NK groups at day seven, even though tumor radiance differences were not yet significant (**Figure S6**). Tumor loads was excluded for time point "D28" due to cell count scarcity. The model included random intercept effects for time and group to consider temporal and inter-group variations. $M_{nk}$ was defined as:

$$\mathbf{Y} = \mathbf{X_{NK}\beta_{NK}} + \mathbf{Z_{NK}u_{NK}} + \boldsymbol{\epsilon}_{\mathbf{NK}},$$

where $\mathbf{Y_1} = (y_k)_K$: mean average tumor radiance in unit of $p/s/cm^2/sr$ of mice in the k-th group at the next time point, $\mathbf{X} = \left(x_{kig}\right)_{K*I*G}$: gene expression of the g-th gene of the i-th cell in the k-th group, $\boldsymbol{\beta} = \left(\beta_g\right)_G$: fixed effects regression coefficient for the g-th gene, $\mathbf{Z} = \left(\mathbf{Z_j}\right)_J$: random effects for time and group, $\mathbf{Z_j} = (t, k)$, $\mathbf{u} = \left(u_j\right)_J$: random effects regression coefficient for the j-th random effect, $u_j \sim N(0, G_{u_j})$ and $G_{u_j}$ is the covariance matrix**,** and $\boldsymbol{\epsilon} = (\epsilon_{ki})_{K*I}$: random error for the i-th cell in the k-th group, $\epsilon_{ki} \sim N(0, \sigma_e^2)$. Fixed effects regression coefficients of significant genes and GOBP pathways were multiplied with -1 to indicate tumor suppressive effect of NK cells and used for subsequent ABM specifications. The fitted significant genes and GOBPs are shown in Table S2 below. The derived coefficients for significant genes and GOBP pathways were incorporated into the modeling of NK cell cytotoxicity using a link function (**Methods**).

Table S2 Significant genes and pathways associated with NK cell anti-tumoral capacity.

| **Covariate** | **EST** | **LL** | **UL** |
|---|---|---|---|
| CD226 | -2.800 | -5.241 | -0.360 |
| NFKBIA | -6.037 | -10.634 | -1.439 |
| SOCS3 | 4.112 | 1.691 | 6.533 |
| RGS2 | 3.893 | 0.989 | 6.797 |
| ZFP36 | -7.017 | -12.141 | -1.894 |
| GZMH | -7.777 | -11.257 | -4.297 |
| GZMK | 4.144 | 1.140 | 7.147 |
| LAG3 | 6.780 | 4.119 | 9.442 |
| PDCD1 | 5.098 | 2.123 | 8.072 |

| | | | |
|---|---|---|---|
| GOBP_NATURAL_KILLER_CELL_MEDIATED_IMMUNITY | 11334.442 | 1922.993 | 20745.890 |
| GOBP_NEGATIVE_REGULATION_OF_NATURAL_KILLER_CELL_MEDIATED_IMMUNITY | 4085.202 | 1464.114 | 6706.290 |
| GOBP_NATURAL_KILLER_CELL_CHEMOTAXIS | -2467.813 | -3757.354 | -1178.272 |
| GOBP_NATURAL_KILLER_CELL_DIFFERENTIATION | -10313.178 | -16955.513 | -3670.843 |

To understand how molecular features drive NK cell cytotoxicity and contribute to tumor control, we applied feature selection to paired scRNA-seq and tumor radiance datasets from lymphoma mice model[5] (**Methods**) and identified gene and GO Biological Process (GOBP) pathway signatures (**Figure 2a**). PDCD1, LAG3, SOCS3, GZMK, RGS2, and gene sets for NK cell-mediated immunity (GO:0002228) and negative regulation of NK cell-mediated immunity (GO:0002716) were positively associated with tumor loads (p-adj < 0.05). PDCD1, the Programmed Cell Death 1 gene, encodes the PD-1 protein that down-regulates NK cell anti-tumor activity[16]. LAG3 is an immune checkpoint receptor that inhibits NK cell immunity[27]. Upregulation of SOCS3 inhibits NK cell terminal maturation and desensitizes NK cells to IL15 stimulation, reducing its antitumor and antiviral activity[28]. RGS2 is a regulator of G protein signaling, which plays a role in modulating NK cell cytotoxicity through the second messenger cAMP[29]. Interestingly, GZMK and GO:0002228 were found to be positively associated with tumor loads, contradicting with their conventional role in NK cell effector function[30]. Tang et al.[22] showed that GZMK were exclusively expressed in $CD56^{bright}CD16^{lo}$ NK cells, which were responsible for immunomodulation and cytokine production in contrary to their cytotoxic $CD56^{dim}CD16^{hi}$ siblings. GO:0002228 contains a repertoire of 80 genes that govern a range of functions from cytotoxicity, cell adhesion, MHC-I recognition, to exhaustion with the inclusion of genes such as FCGR3A, GZMB, and LAG3, presenting potential confounding effects. In addition, the magnitude of their effects was relatively small (GZMK: mean: 1.767 [0.486, 3.047]; GO:0002228: mean: 3.349 [0.568, 6.131]).  On the other hand, CD226, NFKBIA, GZMH, ZFP36, and gene sets for NK cell differentiation (GO:0001779) and chemotaxis (GO:0035747), were negatively associated with tumor loads (p-adj < 0.05). CD226 is a known activating receptor for NK cell cytotoxicity [31], and GZMH directly indicates NK cell effector function[22]. NFKBIA and ZFP36 are stress genes, and NKFBIA was differentially expressed in a cytotoxic $CD56^{dim}CD16^{hi}$ NK subset that was highly inflammatory and exhibited immune-recruiting signatures in Tang et al.[22]. Even using the same NK cell engineering strategies, cells can possess varying anti-tumoral ability. The functional heterogeneity is important for recapitulating experimental variations. Therefore, the above gene and GOBP signatures were then integrated in cytotoxic NK cell agents to model variability in cytotoxicity via a link function (**Methods**).

##### 2.2 Tumor cell viability model $M_{TM}$

In addition to $M_{NK}$, we also built an LME model for tumor cells with scRNA expressions of tumor cells and tumor radiance. Tumor cells are intrinsically programmed to proliferate and survive[18]. We focused on 17 GOBPs governing tumor cell proliferation, cell cycle regulation, and apoptosis. We also included HLA-E, HLA-C, HLA-B, HLA-A, HLA-F, HLA-G, HLA-DOA, HLA-DOB for immune recognition[32], BRAF, NRAS, KIT, MAPK2 for uncontrolled cell growth[33], ERBB4, GRIN2A, and GRM3 for tumor progression,

RAC1 and PREX2 for cell motility and metastasis[34,35], HIF1AN, HIF1A-AS1, HIF1A, and HIF1A-AS2 for hypoxia response[36], and VEGFC, VEGFA, VEGFD, and VEGFB for angiogenesis and metastasis[37]. The complete list is provided in **Supplementary File 3**.

To select significant genes and pathways and quantify their effects on tumor viability, we applied a LME model, $M_{TM}$, to paired scRNA-seq data of tumor cells and tumor radiance data from the lymphoma mouse model[5] using R package lme4[26]. In $M_{TM}$, we proposed that tumor viability was associated tumor scRNA-seq data at the current time point. $M_{TM}$ was defined as:

$$\mathbf{Y} = \mathbf{X_{NK}}\boldsymbol{\beta}_{\mathbf{NK}} + \mathbf{Z_{NK}}\mathbf{u_{NK}} + \boldsymbol{\epsilon}_{\mathbf{NK}},$$

where $\mathbf{Y} = (y_k)_K$: mean average tumor radiance in unit of $p/s/cm^2/sr$ of mice in the k-th group, $\mathbf{X} = \left(x_{kig}\right)_{K*I*G}$: gene expression of the g-th gene of the i-th cell in the k-th group, $\boldsymbol{\beta} = \left(\beta_g\right)_G$: fixed effects regression coefficient for the g-th gene, $\mathbf{Z} = \left(Z_j\right)_J$: random effects for time and group, $Z_j = (t, k)$, $\mathbf{u} = \left(u_j\right)_J$: random effects regression coefficient for the j-th random effect, $u_j \sim N(0, G_{u_j})$ and $G_{u_j}$ is the covariance matrix, and $\boldsymbol{\epsilon} = (\epsilon_{ki})_{K*I}$: random error for the i-th cell in the k-th group, $\epsilon_{ki} \sim N(0, \sigma_e^2)$. The fitted significant genes and GOBPs are shown in Table S3 below.

Table S3 Significant genes and pathways associated with tumor cell viability.

| **COVARIATE** | **EST** | **LL** | **UL** |
|---|---|---|---|
| RAC1 | 43.643 | 14.286 | 73.001 |
| GOBP_POSITIVE_REGULATION_OF_MESENCHYMAL_CELL_PROLIFERATION | -76272.936 | -109095.856 | -43450.017 |
| GOBP_POSITIVE_REGULATION_OF_MESENCHYMAL_STEM_CELL_PROLIFERATION | -45146.802 | -66070.674 | -24222.930 |
| GOBP_REGULATION_OF_CELL_CYCLE_G1_S_PHASE_TRANSITION | 172750.452 | 113223.674 | 232277.230 |
| GOBP_POSITIVE_REGULATION_OF_CELL_CYCLE_G1_S_PHASE_TRANSITION | -66442.154 | -106821.459 | -26062.849 |

We found that RAC1 GOBP for regulation of cell cycle G1-S phase transition (GO:0044843) were positively associated with tumor loads. GOBPs for positive regulation of mesenchymal cell proliferation (GO:0002053), mesenchymal stem cell proliferation, and positive regulation of G1-S transition (GO:1902808) were negatively associated with tumor loads with smaller magnitude, which could be an adjustment to the large magnitude of GO:0044843 due to their synergistic effects in promoting tumor growth. As the CAR-NK cells were engineered to target CD19 on tumor cells, we manually included CD19 expression in lymphoma ABMACT model.

3 ABM fitting

3.1 Mean squared error (MSE) loss

MSE loss was calculated between the simulated data and observed data using the tumor progression ratios between experimental groups and the Raji control group as $L = MSE(r_{sim}, r_{obs})$, where $r_{sim} = \frac{T_g}{T_{Raji}}, r_{obs} = \frac{V_g}{V_{Raji}}, T(t) = \frac{N(t)}{N_0}$ is tumor progression ratio of time $t$, $V$ is the average tumor radiance in $\mathrm{p/s/cm^2/sr}$. As tumor sizes were not directly measured, the normalization against Raji control reduced fitting errors due to scaling discrepancies to allow comparison across models. Data points after post-engraftment day 14 (equivalent of post-infusion day 21) were removed due to lack of data points for the Raji control group.

3.2 Stability

We evaluated simulation stability using coefficient of variation ($CV = \sigma/\mu$) on fitted simulation results of the lymphoma model (**Figure S7**). The results are shown below in Table S4. 30 runs were performed for each experiment group over 35 simulation days. Accumulated tumor progression was calculated for each experiment group and simulation run. Mean and standard deviation were computed by experiment group. While the CD19IL15 CAR-NK had the highest CV, it's likely due to small mean tumor progression. The other two groups had both higher standard deviation ($\sigma$) and higher mean ($\mu$) of tumor progression.

Table S4 Coefficient of variation on fitted simulation results of the lymphoma model

| | $\mu$ | $\sigma$ | CV |
|---|---|---|---|
| CD19IL15 CAR-NK | 27.286 | 7.019 | 25.7% |
| CD19 CAR-NK | 1043.683 | 46.979 | 4.5% |
| NT-NK | 347.520 | 35.730 | 10.2% |

4 Sensitivity Analysis

Aggregating simulation data from in silico perturbation experiments of ABMACT, we trained a Random Forest Regressor (RFR) using scikit-learn[38] to evaluate the importance of model parameters on accumulated tumor growth and prediction accuracy. To quantify the potential gain on time to tumor clearance, we kept other parameters constant and only perturbed the top important features in the CD19IL15 CAR-NK simulations. Tumor growth was measured by the area under the curve (AUC) over a 35-day simulation period. Prediction accuracy was measured by MSE between simulated data and observed experiment data in the xenograft lymphoma mice model in [5]. Feature importance was evaluated by the mean decrease in impurity (MDI) and permutation importance (PI). The top important features of tumor growth AUC measured by MDI and PI are shown in **Figure 5a** and **Figure S5d**. The top important features of model MSE measured by MDI and PI are shown in **Figure S5f-g**. In terms of motility, there is a continuous trend for improved killing with increasing motility (**Figure S5d**), which is expected, as NK cells search and engage tumor cells to exert killing, and higher motility accelerates this process. The top three splitting of model parameters are shown in **Figure S8**.

5 Ordinary differential equations modeling tumor-NK cell dynamics

We compared ABMACT with ordinary differential equations (ODE) modeling and evaluated the performance on the lymphoma mouse model dataset[5]. Adapted from the approach in Kirouac et al.[8], the tumor-NK cell dynamics are described as follows:

$$\frac{dB}{dt} = \mu_B B - p_{k_g} N_c$$

$$\frac{dN_c}{dt} = \mu_g \left( \frac{B^{m_1}}{B50^{m_1} + B^{m_1}} \right) N_c - d_g N_c - \left( 1 - \frac{s_g}{S_g} \right) N_c - p_v \left( \frac{1}{1 + B^{m_2}} \right) N_c$$

$$\frac{ds_g}{dt} = -p_{k_g} + \mu_g \left( \frac{B^{m_1}}{B50^{m_1} + B^{m_1}} \right) S_g$$

$$\frac{dN_e}{dt} = \left( 1 - \frac{s_g}{S_g} \right) N_c - d_g N_e$$

$$\frac{dN_v}{dt} = p_v \left( \frac{1}{1 + B^{m_2}} \right) N_c - d_v N_v$$

Here, tumor cell population $B$ had proliferation rate $\mu_B$. Cytotoxic NK cells $N_c$ could kill tumor cells at a probability of $p_{k_g}$. Different NK cell products possess varying properties, the subscript $g$ denotes parameters specific to NK cell group $g$. $N_c$ population proliferated at a baseline proliferation rate $\mu_g$ and modified by the presence of tumor antigen. The modification effect of tumor antigen was represented by the Hill's function $\frac{B^{m_1}}{B50^{m_1}+B^{m_1}}$, where $B50$ was the half-maximum population count of tumor cells and $m_1$ was Hill's exponent. $N_c$ died at a probability of $d_g$. The exhaustion of cytotoxic NK cells was modelled by the reduction of average serial killing capacity of the population as $\left(1 - \frac{s_g}{S_g}\right) N_c$, where $s_g$ measured the average serial killing capacity of $N_c$, and $S_g$ was the maximum serial killing capacity. $s_g$ was reduce by the average killing rate $p_{k_g}$ and increased by the generation of new cytotoxic NK cells. When tumor cells were cleared, $N_c$ could also transform to the vigilant phenotype $N_v$ at a probability $p_v$. In the modifier $\frac{1}{1+B^{m_2}}$, $m_2$ was a large enough constant so that the transformation only occurred after B approached 0. $N_e$ denoted the number of exhausted NK cells. $d_v$ was vigilant NK cells' death rate. We varied initialization and optimizers to minimize the mean squared error of fitted ratios between experimental groups and Raji control and the observed ratios. The optimal results are provided in **Table S4**.

Table S4 ODE fitting results for lymphoma mouse model

| **group** | **Raji+CD19IL15-CAR NK** | **Raji+CD19-CAR NK** | **Raji+NT NK** |
|---|---|---|---|
| $p_{k_g}$ | 0.013 | 0.006 | 0.010 |
| $\mu_g$ | 0.201 | 0.010 | 0.150 |
| $d_g$ | 0.407 | 0.250 | 0.150 |
| $S_g$ | 20.5 | 20.0 | 15 |
| $p_v$ | 0.924 | 0.900 | 0.900 |
| $d_v$ | 0.001 | 0.001 | 0.001 |
| $B_{50}$ | 25022.880 | 10000.069 | 10000 |
| $m_1$ | 1 | 1 | 1 |
| $m_2$ | 10000 | 10000 | 10000 |
| **optimizer** | Least Squares | Least Squares | LBFGSB |
| **AIC** | 124.020 | 161.811 | 160.262 |
| **BIC** | 127.046 | 164.837 | 163.288 |
| **Train MSE** | 0.013 | 0.028 | 0.024 |
| **Test MSE** | 0.003 | 0.189 | 0.069 |

Table S5 Train and test MSE using ABMACT and ODE

| | **MSE** | **Raji+CD19IL15-CAR NK** | **Raji+CD19-CAR NK** | **Raji+NT NK** |
|---|---|---|---|---|
| **ABMACT** | Train | 0.007 | 0.029 | 0.022 |
| | Test | 0.003 | 0.357 | 0.043 |
| **ODE** | Train | 0.013 | 0.028 | 0.024 |
| | Test | 0.003 | 0.189 | 0.069 |

The train and test MSE in CD19IL15 and NT groups were lower in ABMACT than ODE, although higher MSE were shown in the CD19 group (**Figure S4c, Table S5**). The lower testing accuracy in ABMACT is due to deviations of simulated prediction from tumor outgrowth at day 14 post-engraftment in the CD19 CAR-NK group. We considered reducing cellular properties of CD19 CAR-NK cell agents, such as lower baseline proliferation and killing rates. However, it resulted in sufficient tumor control at day seven post-infusion. The evident increase of tumor radiance in the CD19 group prompted us to speculate imaging artifacts or intrinsic differences in tumor cells in the CD19 group, which could contribute to the outgrowth after day seven post-infusion.

Despite achieving lower losses, the ODE model was limited by its reliance on high-level simplifications. As complexity increased, the model exhibited instability during parameter fitting, especially when dataset sizes were small. Parameters in the ODE model, such as killing rate and proliferation, did not consider the proximity required in cell-cell interactions and dependence on local tumor density and cytokine levels. These limitations reduced the ability of the ODE model to accurately recapitulate immune-tumor dynamics in a biologically meaningful way. Considering both accuracy and interpretability, ABMACT offers advantages by balancing predictive accuracy and mechanistic interpretability. While achieving comparable or superior performance in most groups, ABMACT provides a more detailed and biologically faithful representation of immune-tumor interactions. By decomposing the NK cell lysis of target cells into chimeric antigen recognition, the combination of activating and inhibitory signaling, and NK cell intrinsic properties including baseline cytotoxicity and serial killing capacity, we attempted to provide a mechanistic explanation of the superior tumor control in the CD19IL15-CARNK group versus its experiment control and null control.

#### 6 Software

The genetic and cell functional modeling were performed in R (v.4.2.2)[39] and Python (v3.9)[40] using Seurat [41], lme4[42], scipy[12]. The ABM was built in Python using MESA[1].
